# Computational design of allosteric pathways reprograms ligand-selective GPCR signaling

**DOI:** 10.1101/2025.08.13.670154

**Authors:** M. Hijazi, D. Keri, A. Oggier, A. Sengar, M. Agosto, T. Wensel, P. Barth

**Affiliations:** Institute of Bioengineering, School of life sciences, EPFL, Lausanne, Switzerland; Verna and Marrs McLean Department of Biochemistry, Baylor College of Medicine, Houston, TX, USA

**Author notes:** equal contribution (M. Hijazi, D. Keri, A. Oggier).

## Abstract

G-protein-coupled receptors (GPCRs) constitute the largest family of signaling receptors and drug targets. However, understanding how variations in receptor sequence, ligand chemical structure, and binding impact signaling functions remains a challenge, hindering drug discovery. Here, we developed a computational protein structure and dynamics approach to infer and design GPCR responses to multiple ligands. We created 32 dopamine D1 and D2 receptor variants with widely reprogrammed agonist-induced signal transductions. Subtle natural and designed receptor sequence variations, predicted to alter specific structural and dynamic mechanisms of ligand responses, profoundly impacted ligand potency and efficacy in agreement with our calculations. Our study provides a rational blueprint for computing the effect of sequence polymorphisms on ligand-selective protein signaling and paves the way for advancements in pharmacogenomics, drug selectivity, and the design of signaling receptors from first principles.

## Introduction

Signal transduction is a universal mechanism of communication in living cells, controlling a wide range of physiological processes. Specialized proteins, including GPCRs, carry out this process by translating extracellular stimuli into specific intracellular functions. Due to their critical roles in numerous diseases, GPCRs have become one of the largest classes of drug targets. However, many drug compounds suffer from off-target effects, lack of intracellular pathway selectivity, and strong pharmacological sensitivity to genetic variations in the human genome^1–3^.

Structure-function studies have revealed that ligands trigger precise intracellular responses by contacting selective subsets of residues in the receptor binding site^4^. Hence, functional mapping of GPCR binding surfaces should accelerate the rational design of intracellular pathway-selective drugs. However, these studies currently rely on extensive experimental characterization of receptor-ligand systems^5,6^. Natural genetic variation in human GPCRs leads to individual differences in responses to medication^2^. For example, subtle sequence alterations at locations distant from the binding sites can profoundly affect signaling responses to drugs^2,4,7^, implying long-range (i.e., allosteric) control of signal transduction by receptor structure and dynamic properties. Predicting drug efficacy and functional selectivity as a function of receptor polymorphism would therefore be highly desirable, but it requires a quantitative understanding of GPCR allosteric communications that has remained elusive so far.

Current computational techniques either predict the structure and binding of conformationally stable proteins^8,9^, or study the conformational dynamics of receptors often in the absence of binding partners^10,11^. However, these approaches cannot predict and design ligand-mediated receptor allosteric signal transductions. Here, we developed a computational framework that combines molecular dynamics, information theory, protein docking, and design to understand how receptor sequences, structures, and dynamics encode selective responses to ligand chemical structures and binding modes. Using this framework, we uncovered the molecular mechanistic underpinnings of ligand-mediated GPCR signaling in the dopamine receptor family, which forms critical nodes in the pathways targeted by most current anti-psychotic drugs.

We leveraged this knowledge to predict the impact of sequence variations on drug responses and design a wide range of receptors with high ligand selectivity. Our findings and methods pave the way toward mapping functional sites and signal transduction pathways in GPCRs, as well as informing the rational design of personalized and selective drugs.

### Rationale for the prediction of ligand-selective signaling responses

GPCRs are highly flexible molecules that can be regulated by multiple ligands through the selection of specific receptor conformations^3,15–18^. For example, agonist ligand binding to the receptor extracellular surface triggers cell signaling by stabilizing receptor conformations on the intracellular side that are prone to G-protein or beta-arrestin recruitment and activation. Such long-distance communication can be achieved by networks of residues that efficiently propagate ligand-induced changes in receptor structure and dynamics across the protein scaffold^19–22^. By mechanically coupling distant sites on the protein surface, these networks may enable allosteric regulation of receptor function. While chemically similar ligands can regulate unrelated proteins, ligand agonists with different chemical structures and binding modes can activate the same receptor^23,24,25^. How, then, can diverse chemical inputs propagate into selective cellular responses through the same protein scaffold? We hypothesize that receptors have evolved to encode various allosteric pathways engaged by distinct ligand chemical groups to communicate with the receptor intracellular surfaces (**Fig.1a, b**). This multiple pathway view of intramolecular signal propagation offers a conceptual framework for predicting, testing and understanding the relationships between receptor-ligand sequence-structure properties and signaling functions. To interrogate this model, we selected two extensively characterized and evolutionarily distant members of the dopamine receptor family, the D1 and D2 receptors. While being primarily activated by the same ligand agonist, dopamine, these receptors share only 19% sequence identity and activate two distinct G-protein pathways, Gs for D1 and Gi for D2. As such, the D1- D2 pair constitutes a paradigm for studying the relationships between the receptor sequence-structure context and activation mediated by the same ligands.

**Figure 1.**
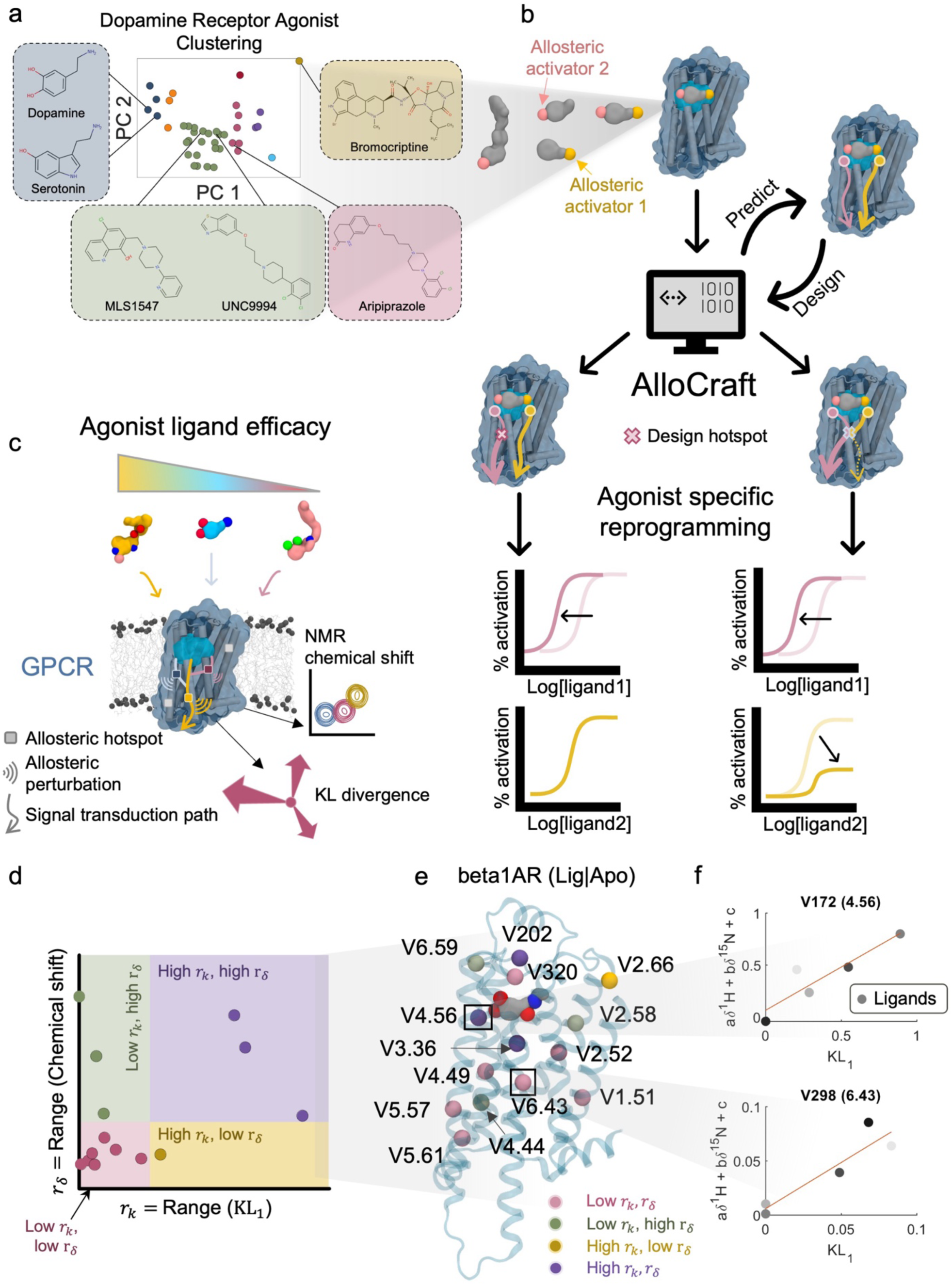
Hypothesis and approach for assessing GPCR’s response to multiple ligands. **a.** Principal component analysis plot of dopamine receptor agonists clustered according to chemical structure similarity. Representative agonist structures of each cluster are highlighted. **b.** Computational and experimental workflow for receptor design. Upon selection of agonists, molecular dynamics (MD) simulations of ligand-bound receptors are carried out and analyzed by the software AlloCraft to predict ligand-specific allosteric pathways. Allosteric hotspots are targeted for in silico mutagenesis to design receptors with ligand-selective responses (**Fig.S1, Fig.S2**). **c.** Ligands with distinct allosteric efficacy engage different allosteric transduction pathways reaching the intracellular receptor surface. Structural perturbations at specific hotspots along the allosteric transmission pathways can be identified experimentally via NMR spectroscopy or computationally via Kullback-Leibler divergences (Kldiv) from MD simulations**. d.** Classes of structural perturbations upon ligand binding measured as ranges of NMR chemical shifts (*r*_*δ*_) or ranges of KLdiv (*r_k_*) for the β1AR YY variant. **e.** Beta1AR structure (transparent cartoon) with the valine sites reported in Grahl et al.^26^ highlighted according to their *r_k_*, *r*_*δ*_ classification. **f.** Examples of two valine sites, V4.56 and V6.43, showing significant correlation between *KL*_#_ and an NMR chemical shift linear fit (*aδ*^1^*H* + *bδ*^15^*N* + *c*) for agonists with different efficacies (V4.56: *R*^2^ = 0.831; V6.43: *R*^2^ = 0.852). Full comparison between AlloCraft predictions and NMR measurements are reported in **Fig.S3-Fig.S7**.

We first analyzed the structural space of known ligand agonists of the D2 receptor based on their chemical structure similarity. When mapped along the first 2 principal components, agonists clustered into several families of structures that primarily differ in size and chemical groups (**Fig.1a**). Each chemical group can, in principle, define a different allosteric activator moiety that binds specific receptor functional sites and engages selective activation pathways (**Fig.1b**). If this hypothesis is valid, we should be able to rationally design receptors with altered, highly selective responses to specific ligands by introducing sequence alterations that rewire the residue networks defining each pathway.

To address this question, we developed the computational prediction and design method AlloCraft. AlloCraft infers receptor intramolecular communication residue networks from atomistic molecular dynamics simulations of GPCR structures and information theory-based interpretation of protein structure dynamics (**Methods, Fig.1b, Fig.S1, Fig.S2**). The method identifies two forms of allosteric perturbations: structural changes, measured as shifts in dihedral angles via Kullback-Leibler (KL) divergence, and dynamic changes, measured as coupled motions between residues via Mutual Information (MI). AlloCraft then tests these pathway predictions through sequence-structure perturbations designed to mimic natural genetic variations and reprogram selective ligand-mediated allosteric responses.

### Prediction of allosteric pathways inferred from NMR spectroscopy

Before analyzing the dopamine receptor family, we first evaluated whether our method could accurately predict allosteric signal transduction pathways in GPCRs. The β1-adrenergic receptor (β1AR) was chosen as a benchmark, given that structural perturbations caused by sequence variations, ligand binding, and intracellular protein interactions have been extensively characterized using NMR spectroscopy^26,27^. These studies tracked structural changes through chemical shift (CS) variations in isotopically labeled amino acids, revealing long-range communication between the ligand-binding site and distal regions of the receptor. Although CS variations do not directly measure residue dynamics or transitions between conformational states, they do provide evidence for allosteric interactions—namely, coupling between ligand-binding residues and distant sites within specific receptor conformations—and how these interactions shift upon binding to different ligands. This makes CS data well-suited for comparison with AlloCraft’s KL divergence-based ‘structural allostery’ metric (**Fig.S1**), which quantifies structural changes between different ligand-bound states of a receptor (**Fig. 1c**).

NMR measurements on β1AR labeled with isotopically labelled valines revealed that the signals of several valine residues—positioned far from the ligand binding site—strongly correlated with ligand efficacy, indicating robust long-range allosteric communication. To test whether AlloCraft could recapitulate these allosteric effects, we modeled and simulated 10 β1AR-ligand complexes across different functional states and evaluated allosteric responses of 28 valine positions to 5 ligands in two receptor sequence variants (**Fig. 1d–f**, **Fig.S3**). One variant was a thermostabilized (TS) version of β1AR, while the other (YY) was closer to the wild-type (WT) sequence, preserving two native tyrosine residues essential for activation and G-protein coupling^26^.

Allosteric perturbations from ligand binding or sequence mutations were quantified using Kullback-Leibler divergence (KLdiv)^28^, which measures conformational differences across ensembles. We also calculated Mutual Information (MI) to assess dynamic couplings between the ligand binding site and specific residues (Methods). Our KLdiv predictions showed good qualitative agreement with NMR-based perturbations for 21 out of 28 valines (**Fig. 1d–f**, **Fig.S4**). Except for 3 sites, we also observed strong correlation between the KLdiv and CS values (**Fig.S5**, **Fig.S6**). Of the remaining 7 valine residues, 5 showed no correlation between ligand efficacy and NMR chemical shifts, suggesting that CS variations at these positions do not capture allosteric communication with the ligand binding site. We therefore expected disagreement with our KLdiv- based structural predictions at these sites.

Interestingly, for these 5 sites, we observed a strong correlation between ligand efficacy and dynamic communication, as captured by MI, suggesting that allosteric effects here are more dynamic than structural in nature and therefore not well reflected by CS measurements (**Fig.S7**). Overall, these results indicate that our approach can identify critical allosteric sites mediating signal transmission and recapitulate structural/dynamic perturbations that are sensitive to the ligand chemical structure, binding mode and receptor sequence.

Following this initial validation, we investigated the dopamine receptor-ligand agonist systems using AlloCraft. To directly test the sensitivity of the method and assess whether distinct ligands activate the D2 receptor differently, we first selected dopamine (DA) and bromocriptine (BRC) because these ligands exhibit highly divergent structures but share similar pharmacology^29^ (**Fig.1a**). Consistent with distinct binding modes (**Fig.S8, Fig.S9**), AlloCraft revealed that DA and BRC communicate with the G-protein binding surface through allosteric signal transduction pathways that only partially overlap (**Fig.2a,b**). While both ligands use transmembrane helix 5 (TM5) as a strong communication channel, they each engage distinct networks of allosteric residues running through TMs 3, 4, 6 and 7 (**Methods**).

**Figure 2.**
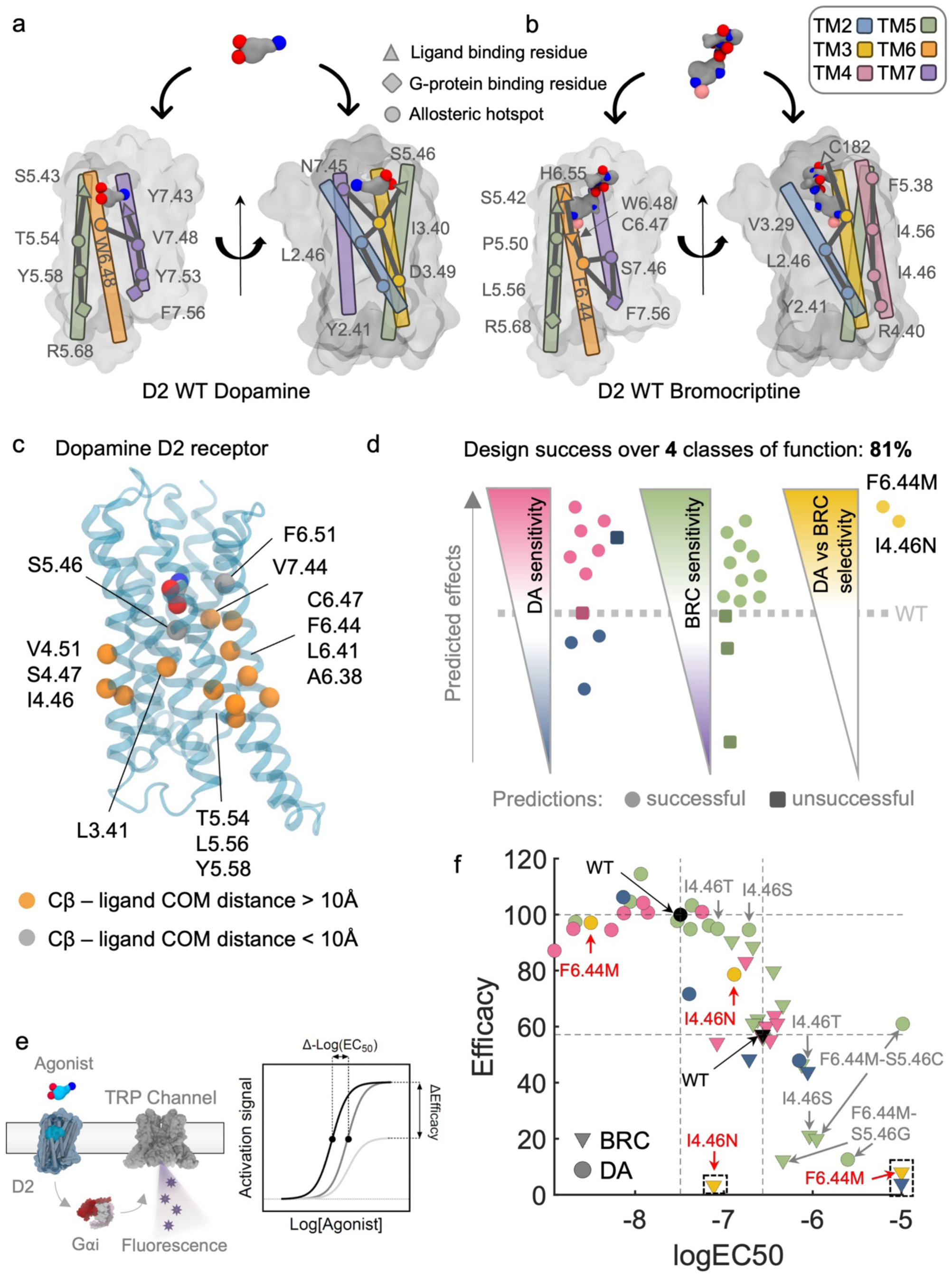
Functional validation of the D2 designs. **a.** and **b.** Highlights of allosteric pathways extracted for dopamine bound (**a.**) and bromocriptine bound (**b.**) dopamine D2 receptor (**Methods**, **Fig.S1-2**) **c.** Positions of the mutated sites shown on the dopamine D2 receptor structure. **d.** Classes of design according to their predicted effect: 1) class I (PINK), II (BLUE) for enhancing or decreasing the sensitivity to DA; 2) class III (GREEN) for enhancing the sensitivity to BRC; 3) class IV (YELLOW) predicted to increase DA selectivity versus BRC. Each point corresponds to a designed receptor and is located on the scales according to the changes in ligand potency from WT. Successful and unsuccessful predictions are displayed using filled circles and black squares, respectively. **e.** Schematic of the TRP channel assay used to characterize D2- mediated Gi activation. Changes in ligand potency and efficacy for each D2 receptor are extracted from dose response curves measured using the assay (**Fig.S11-Fig.S15**). **f.** DA and BRC efficacies and log(EC50) values normalized for receptor cell surface expression measured by ELISA for the D2 variants. The data points are colored according to the predicted functional effect from **c.** Black data points represent the WTs. Boxed data points correspond to variants for which measured signals were too low to quantitatively estimate efficacy and potency. Red labelled variants showed the largest changes in ligand selectivity. Named variants were further investigated.

If these predictions are accurate, we should be able to reprogram specific ligand-mediated D2 responses through sequence alterations at allosteric sites that belong to ligand-selective pathways. Ligand-specific effects may also be obtained by targeting sites engaged by both ligands if designed sequence-structure perturbations rewire paths in each receptor-ligand complex differently. Lastly, the dynamic interactions between allosteric residues can be modulated by their environment. Therefore, residues neighboring these sites represent additional targets for redesigning allosteric signal transduction.

### Computational design of ligand-selective D2 signaling responses

We explored these different scenarios using AlloCraft and scanned the TM region for sequence variations that altered signaling responses. This design perturbation approach was followed by experimental validation to provide a stringent validation of our allosteric pathway predictions.

The method selected amino-acid mutations (among 20 possible substitutions at each site) by first assessing their impact on the equilibrium between inactive and active state conformations (**Fig.S1**). Mutations predicted to cause significant changes in conformational stability—leading to receptor inactivation, unfolding, or high constitutive activity—were therefore discarded from further consideration. Remaining mutations were ranked based on their impact on correlated motions of allosteric sites, which we calculate and use as a proxy for allosteric coupling and signal transduction capacity^7,29^. Sequence variations increasing the allosteric coupling in the ligand and G-protein-bound active state complex should enhance ligand-mediated G-protein activation and vice versa. Overall, AlloCraft scanned a total of 39 positions and created 26 D2 variants with 4 classes of predicted functional effects through designed mutations at 14 positions located on all TMs except TMs 1 and 2 (**Fig.2c,d**). Class I and II aimed to enhance or decrease the sensitivity to DA respectively, while class III was designed to enhance sensitivity to BRC. Class IV sought to increase selectivity for DA versus BRC. While classes I, II and III considered the impact of designed mutations on the signal communication of one ligand only (DA or BRC), variants in class IV were obtained by calculating the allosteric coupling for both ligands and maximizing the difference between those quantities.

Since all designed mutations (with the exception of 5.46 and 6.51) lie far away from the ligand and G-protein binding surfaces (i.e. distant from at least 12 Angstrom, **Fig.2c, Fig.S10**), they should not directly perturb binding but instead, regulate signal transduction through long-range allosteric effects. We experimentally characterized ligand-induced Gi activation mediated by these receptors using mammalian cell signaling reporter assays (**Fig.2e**, **Methods**). We extracted ligand potencies (EC50) and efficacies for all receptors from ligand dose responses and then analyzed the impact of the designed mutations on each compound (**Fig.2f**, **Fig.S11-Fig.S15**). Overall, the experimentally measured ligand-mediated signaling activities agreed well with our computational predictions (**Fig.2d**), with 81% of the designed variants achieving the intended DA or BRC ligand selective effects. In particular, the DA-selective class IV variants were correctly predicted and considerably lost sensitivity to BRC while maintaining significant DA-mediated activity (**Fig.2d,f)**.

### Mechanistic underpinnings of reprogrammed D2 signaling

The ensemble of designed D2-ligand systems with widely reprogrammed agonist-mediated activities and selectivity offers a unique opportunity for exploring the mechanistic underpinnings of signal transduction perturbations. We selected representative receptor variants with distinct selectivity and activity for in-depth mechanistic investigation of structural and allosteric dynamic properties. We specifically sought to understand how the single point mutations I4.46N and F6.44M could achieve such high selectivity for DA.

Position 4.46 is located on the intracellular half of the TM4 far away from the ligand binding site and constitutes a BRC-selective allosteric propagation site (**Fig.2a,b**). It belongs to a network of allosteric residues forming a path engaged by BRC but not by DA that runs from the extracellular side of TM5 and propagates through TM4 down to the G-protein binding surface.

Our analysis of the D2–BRC complex revealed that I4.46N functions as an allosteric sink, inducing substantial structural and dynamic changes within the receptor that prevent signal propagation to the G-protein binding interface, consistent with the observed loss of function (**Fig.3a-b, Fig.S16**). Through the formation of new polar contacts, N4.46 triggers conformational reorientation of the neighboring Y2.41, S2.45 and L2.46 side-chains (**Fig.3c, Fig.S16**).

**Figure 3.**
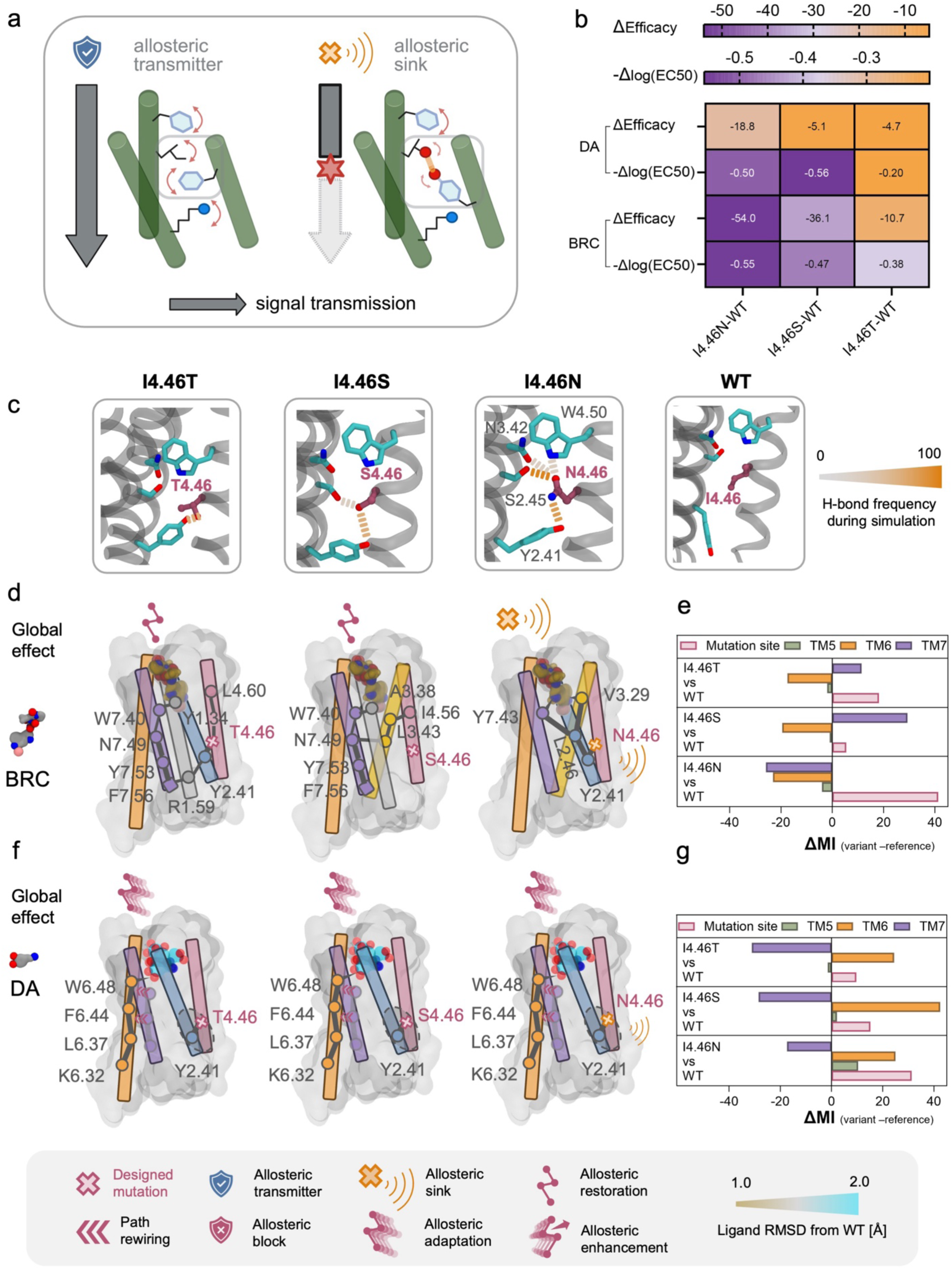
A conformational lock acts as an allosteric sink blocking Bromocriptine-mediated signaling. **a.** Schematic description of the impact of a conformational lock on signal transduction. While weak interactions between side-chains enable correlated motions and signal propagation (left), strong interactions lock side-chain conformations can block signal transduction by breaking the chain of coupled motions. **b.** Changes in ligand efficacy and potency from D2 WT of designed mutations at site I4.46. **c.** Representative snapshots of MD simulations describing the local interaction networks around position 4.46. The strength of polar interactions is derived from their frequency during the simulations. **d, f.** Schematic representation of the main effects of I4.46 mutations on the ligand-specific allosteric pathways calculated by AlloCraft (bromocriptine: **d**; dopamine: **f**). Ligands are colored according to the changes in their binding conformation from their respective position in WT as measured by RMSD over all heteroatoms. The legend describes the allosteric effects observed in the simulations. **e, g.** Mutational impacts on the Mutual Information computed by AlloCraft for key allosteric hub residues forming the principal allosteric pipelines connecting the ligand-binding site to intracellular signaling surfaces (**Fig.S1-2**). ΔMI values are reported for the mutation site and for the pathways running through TM5, 6 and 7 (bromocriptine: **e**; dopamine: **g**). For each pathway, the reported value corresponds to the cumulative ΔMI across the principal allosteric hubs that constitute the top three allosteric pipelines (**Methods**). Positive delta MI values are interpreted as an increase in allosteric communication and vice versa. BRC: While ΔMI values for I4.46N are negative across all TMs, consistent with the observed loss of signaling, strong allosteric communication is restored along TM7 in I4.46S and I4.46T. For DA, compared to WT, the ΔMI variations across TMs largely offset one another in all three variants, aligning with the minimal impact of these designs on DA-mediated signaling.

The I4.46N substitution also allosterically modifies the conformation of ECL3, G4.63 and sidechain of T7.39 that are in close contacts with BRC (**Fig.S16**). These structural changes strengthen the participation of L2.46 and Y2.41 in signal propagation at the junction of allosteric pathways connecting ligand binding regions (through V3.29, F6.51, and Y7.43) to TM2 and the mutation site on TM4. This structural reorganization propagates information preferentially through TM2, reducing the dynamic communication between G-protein binding regions and the rest of the receptor (**Fig.3d-e, Fig.S16**). Overall, I4.46N acts on BRC as an “allosteric sink”, diverting signal propagation away from the effector binding surface toward the mutation neighborhood and reducing overall BRC’s capacity to trigger G-protein activation (**Fig.3d-e**).

We next examined how DA overcomes this mutation and maintains significant Gi signaling efficacy (**Fig.3b**). Unlike for BRC, the structural effects of I4.46N in the D2 receptor were very localized, mostly affecting Y2.41 and did not result in significant signal propagation funneling through TM2 (**Fig.3f-g, Fig.S17**). Instead, we observed significant structural changes in the ligand binding site and conformational adaptation of DA which shifts binding pose in response to the mutation (**Fig.S17, Fig.S18**). Consequently, W6.48, through enhanced ligand contact, replaced Y7.43 to become a major allosteric propagation site for DA. Since this change led to a shift in allosteric communication from TM7 to TM6 (**Fig.S17**), we conclude that I4.46N acts on DA through “allosteric rewiring” (**Fig.3f**).

To further validate AlloCraft’s mechanistic and structural underpinnings of these allosteric effects, we conducted additional mutagenesis at position 4.46. Based on our hypothesis that conformational reorientation through formation of a polar interaction network in this region drives the allosteric sink, we tested whether modulating the polarity and strength of N4.46’s polar contacts with Y2.41 and S2.45 would impact signal transduction. Specifically, we substituted I4.46 with threonine and serine, which were predicted to reduce the number and strength of polar interactions (**Fig.3c**). In our calculations, both substitutions weakened the allosteric sink effect at position 4.46 in the BRC-bound complex and partially restored strong native-like allosteric pathways, primarily along TM7, connecting the ligand-binding site to the G-protein interface (**Fig.3d,e**). Consistent with these predictions, we observed significant BRC-induced signaling for I4.46S and I4.46T, with responses falling between those of I4.46N and WT (**Fig.3b, Fig.S11**) In contrast, these substitutions had minimal impact on dopamine-induced responses, which remained comparable to WT (**Fig.3b**), in agreement with our calculations (**Fig.3f,g**).

These findings validate our AlloCraft analysis and demonstrate that rationally designed polar contacts at position 4.46 can modulate the orientation and conformational dynamics of Y2.41 and S2.45, playing a key role in establishing an allosteric sink.

We next investigated the mechanistic underpinnings of F6.44M’s high DA selectivity. Consistent with the strong loss of function for BRC, our calculations of the WT complexes suggested that F6.44 is a key allosteric transmission site for BRC but not for DA (**Fig.2a**). In the D2-BRC complex, the mutation triggered significant side-chain conformational rearrangements of the neighboring I3.40 and L6.41 which blocked signal propagation through TM6 (**Fig.4**, **Fig.S19**). This effect was very specific to the substitution to methionine as the F6.44I mutant, which did not lose sensitivity for BRC (**Fig.S20, Fig.S21**), displayed strong communication through I6.44 and did not block signal transduction through TM6 (**Fig.3e, Fig.S20**).

**Figure 4.**
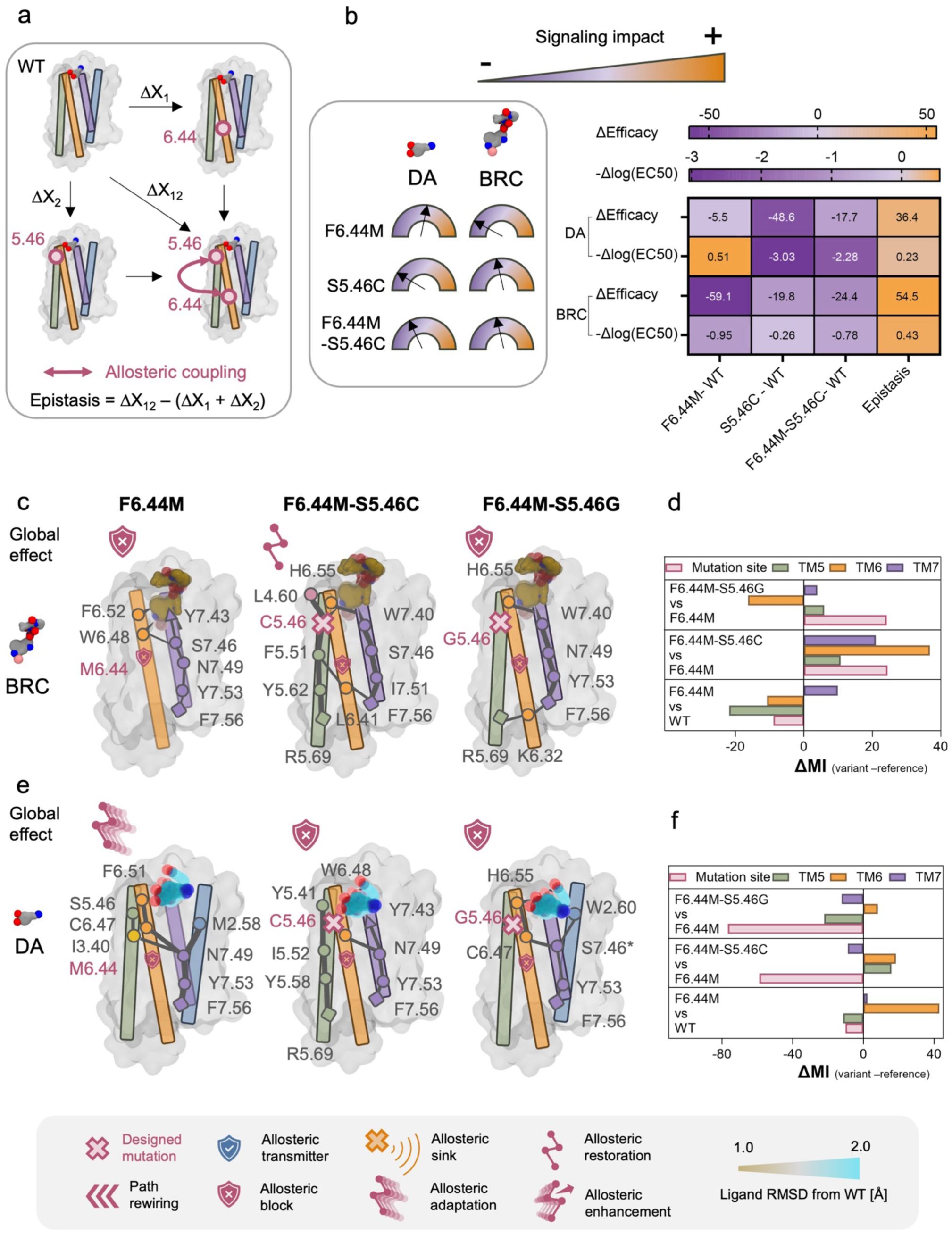
Ligand conformational flexibility enables allosteric rewiring and adaptation. **a.** Thermodynamic description and calculation of epistasis. **b.** Signaling impact of single and double designed mutations at sites 6.44 and 5.46. Schematic description (**left**). Measured changes in efficacy and potency from WT (**right**). Strong epistasis indicates the existence of an allosteric coupling between both sites. **c, e.** Schematic representation of the main effects of designed mutations on the ligand-specific allosteric pathways calculated by AlloCraft (bromocriptine: **c**; dopamine: **e**). Ligands are colored according to the changes in their binding conformation from their respective position in WT as measured by RMSD over all heteroatoms. The legend describes the allosteric effects observed in the simulations. **d, f.** Mutational impacts on the Mutual Information computed by AlloCraft for key allosteric hub residues forming the principal allosteric pipelines connecting the ligand-binding site to intracellular signaling surfaces (**Fig.S1-2**). Delta MI values are reported for the mutation site and for the pathways running through TM5, 6 and 7 (bromocriptine: **d**; dopamine: **f**). For each pathway, the reported value corresponds to the cumulative ΔMI across the principal allosteric hubs that constitute the top three allosteric pipelines (**Methods**). Positive delta MI values are interpreted as an increase in allosteric communication and vice versa. For BRC, ΔMI values for F6.44M are negative along most TMs, consistent with the observed loss of signaling. In contrast, strong allosteric communication is restored across all TMs in F6.44M-S5.46C relative to F6.44M, and to a lesser extent in F6.44M-S5.46G, mirroring experimental measurements. For DA, compared to WT, ΔMI changes are dominated by a positive shift along TM6 in F6.44M, consistent with its gain-of-function profile. Relative to F6.44M, ΔMI variations in F6.44M-S5.46C largely cancel out across TMs—except at the mutation site— correlating with its minimal impact on DA efficacy. In contrast, F6.44M-S5.46G shows predominantly negative ΔMI shifts across TMs, in line with the observed loss of function.

In the context of the DA-D2 complex, F6.44M triggered smaller structural changes in the receptor core than in the ligand binding site (**Fig.S22**). DA shifted binding pose and established stronger binding and allosteric contacts with TM5 (S5.46), TM6, (F6.51), and TM7 (Y7.43) (**Fig.S22**, **Fig.S8, Fig.S9**), consistent with the enhanced measured ligand potency. While M6.44 still acts as an allosteric block, DA repositioning in the binding pocket enables signal transmission to reach the G-protein binding surface through novel communication connections between TM3, TM6 and TM7 (**Fig.3g**).

To further validate that DA establishes stronger allosteric contacts with S5.46, F6.51 and Y7.43, we performed extensive *in silico* mutagenesis at the three target sites to identify substitutions that could allosterically modulate the effect of F6.44M on signal transduction.

Given its central role in orienting D3.32 and interacting with the amine groups of ligands^50^, all Y7.43 mutations were predicted to destabilize the binding site conformation and disrupt ligand binding. Consistent with these predictions, none of the Y7.43 variants exhibited significant BRC- induced signaling activity.

More interestingly, for position F6.51, our calculations indicated that substitution to His could enhance the allosteric coupling of the ligand with the active form of the receptor. In line with these predictions, the F6.51H variants partially rescued the signaling deficits caused by F6.44M on BRC, i.e. improving efficacy and potency, respectively (**Fig.S15**).

The most compelling findings emerged at position S5.46, a known binding hotspot for both DA and BRC (**Fig.S8, Fig.S9**). Although mutations at this site were expected to alter ligand binding affinity (**Fig.S23**), our models predicted that substitutions to Cys, and to a lesser extent Gly, would also enhance allosteric coupling and restore signal transduction mediated by BRC in the F6.44M background (**Fig. 4, Fig.S14, Fig.S15**). Experimental validation confirmed these predictions: the F6.44M–S5.46C double mutant showed a significant restoration of BRC-mediated signaling, with increases in efficacy and potency of up to 34% and 1.5-fold, respectively (**Fig. 4b**). Similarly, F6.44M–S5.46G improved BRC efficacy by 20% (**Fig.S13- Fig.S15**). Interestingly, while S5.46C enhanced BRC signaling in comparison to F6.44M, it reduced DA response—decreasing efficacy by 12% and potency by 620-fold (**Fig. 4b**)—consistent with our calculations (**Fig. 4e-f, Fig.S13- Fig.S15**).

To probe potential allosteric communication between S5.46 and F6.44, we characterized the single S5.46C mutant and, in combination with previous data for F6.44M, computed the epistasis effect for the double mutant (**Fig. 4b**). The observed effects were clearly non-additive, revealing epistatic gains in efficacy for both ligands (55% for BRC) and a 3-fold increase in BRC potency.

While F6.44M is a gain-of-function mutation for DA and a loss-of-function mutation for BRC, the S5.46C addition reversed these outcomes. Thus, S5.46C acts as a negative allosteric regulator of F6.44M for DA, but a positive allosteric regulator for BRC.

In summary, our results revealed a previously unrecognized allosteric connection between positions 5.46 and 6.44, which was accurately predicted and exploited by AlloCraft to design ligand-specific allosteric effects and reprogram D2 receptor signaling.

Overall, our structural investigations of ligand-specific allosteric effects in the designed D2 receptors indicate that ligand chemical structures and binding flexibility are key for exploiting the plasticity of signal transduction pathways and accommodating sequence alterations. To further validate this mechanistic hypothesis, we investigated the effects of chemical modifications on the ligands. Specifically, we assessed how the I4.46N and F6.44M mutations influenced D2 receptor signaling in response to apomorphine (AP), a dopamine analog and known agonist. While AP bears the same agonist chemical moieties as DA, it is bulkier, displayed limited conformational flexibility in our MD simulations and did not adopt alternative binding poses in response to the mutations (**Fig.S24**). In agreement with our simulations, AP-mediated D2 activation was significantly more perturbed by the designed mutations than with DA (**Fig.S24**). Overall, these results provide compelling evidence that ligand binding flexibility is a key mechanism of structural adaptation to sequence variations impacting allosteric communications.

Overall, this study highlights the power of our method to identify and reprogram ligand-specific allosteric pathways, while disentangling the distinct contributions of receptor sequence and structural variations to ligand efficacy and potency.

### Designed mutational effects on the distant D1 receptor

We next investigated how the overall sequence and structure context of the receptor itself modulate the impact of such allosteric mutations on ligand pharmacology. While D1 and D2 share only 19% sequence identity, DA and BRC activate both receptors and trigger signaling primarily through Gi for D2 and Gs for D1 (**Fig.5a**). Structural analysis of ligand binding to D1 reveals that DA and BRC adopt similar binding poses to those observed in D2 (RMSD within 1.2 Å, **Fig.S8, Fig.S9**, **Fig.S25**). DA primarily activates the receptors by engaging with the conserved Ser 5.42, 5.43, and 5.46 on TM5 and Phe 6.51 and 6.52 on TM6. DA also contacts Y7.43 in D2 and the chemically similar W7.43 in D1; however, in this context, only the Y7.43 interaction is involved in signal propagation. 89% of the binding contacts with BRC are conserved in both receptors. We studied the response of two strong (DA and BRC) agonists to five allosteric mutations in D1 (L3.41H, I4.46N, F6.44I, F6.44M, C6.47L) that significantly impacted ligand sensitivity and selectivity of D2. Ligand-induced D1-mediated activation of the G protein Gs was measured in vitro using HEK reporter cell lines transfected with the EPAC cAMP assay.

Ligand dose titrations revealed very different pharmacological impact of sequence alterations when compared to D2. None of the D1 variants exhibited any loss of function and were either neutral or gain of function for the tested ligands. We also observed significantly increased efficacy of DA while only potency was enhanced with D2. Lastly, both I4.46N and F6.44M variants that had completely lost sensitivity to BRC in D2 either kept strong WT-like response or even gained sensitivity for BRC in D1 (**Fig.5b, Fig.S26, Fig.S27**). Structural and dynamic investigations of D1 I4.46N revealed that, while signal transduction is shifted towards TM4 as in D2, N4.46 is not acting as a sink but strengthens signal propagation to the intracellular surface by establishing strong allosteric connections with F2.41, R55 (ICL1) all the way down to Y7.53 which is a critical G-protein binding site (**Fig. 5c-e**).

**Figure 5.**
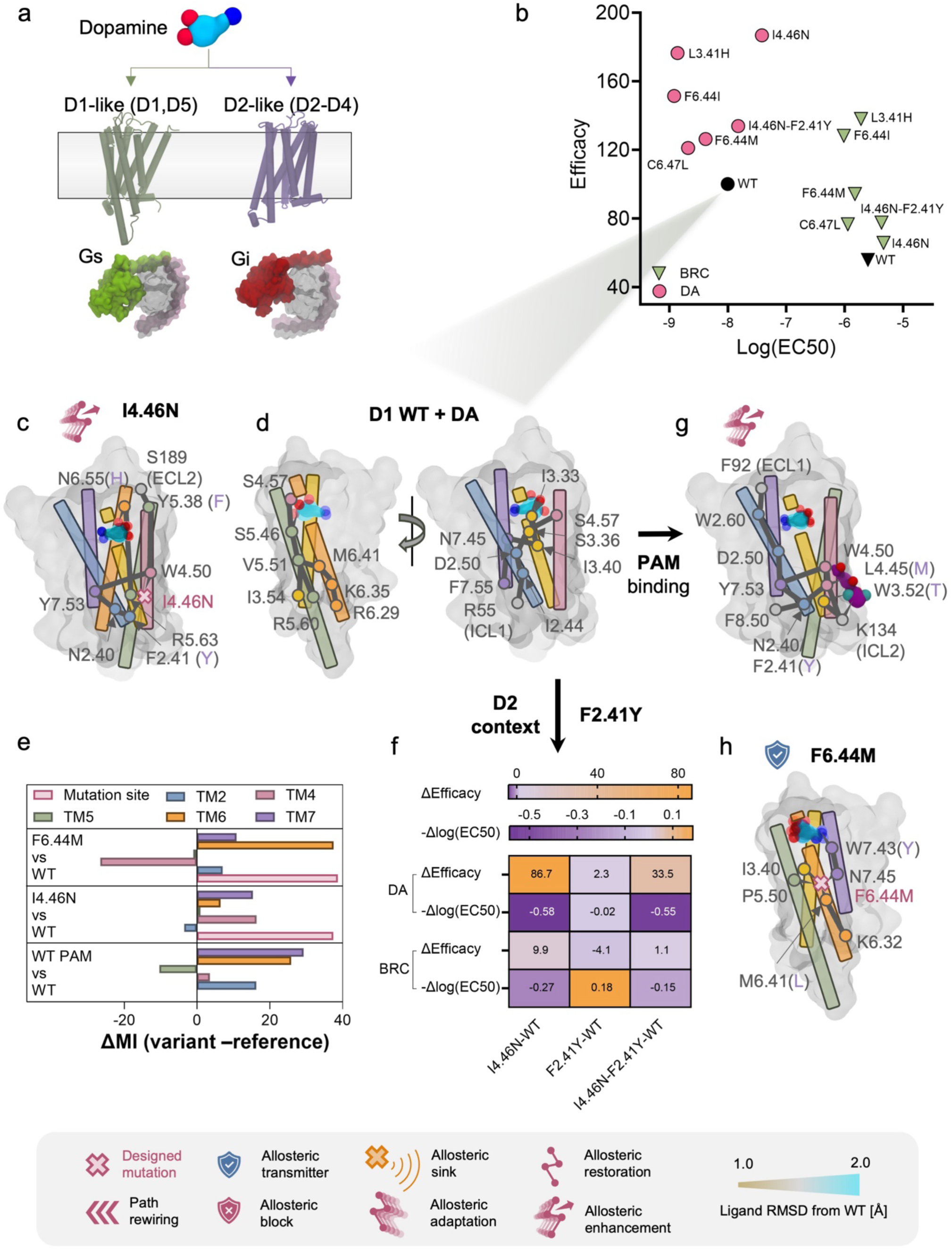
Receptor context greatly impacts the effects of sequence variation on ligand responses. Comparison between dopamine D1 and D2 receptors: **a.** dopamine D1 and D2 receptors are evolutionarily distant (19% sequence identity in humans) and bind to different G-proteins. **b.** Efficacy and potency values of bromocriptine and dopamine for D1 receptors. **c.** Schematic representation of the main calculated allosteric effect of I4.46N in the DA-D1-Gs complex (i.e. major path differences from D1WT). **d.** Schematic representation of the major allosteric pathways for the DA-D1 WT-Gs complex split in 2 views. **e.** Mutational impacts on the Mutual Information computed by AlloCraft for key allosteric hub residues forming the principal allosteric pipelines connecting the ligand-binding site to intracellular signaling surfaces (**Fig.S1- 2**). Delta MI values are reported for the mutation site and for the pathways running through TM2, TM4, TM5, 6 and 7. For each pathway, the reported value corresponds to the cumulative ΔMI across the principal allosteric hubs that constitute the top three allosteric pipelines (**Methods**). Positive delta MI values are interpreted as an increase in allosteric communication and vice versa. Compared to WT, ΔMI changes for each variant are dominated by a positive shift along most TMs, consistent with their gain-of-function profile. **f.** Impact of the D2 residue Y2.41 on D1 signaling response to dopamine. **g.** Allosteric enhancing effect of positive allosteric modulator (PAM) LY3154207 binding to D1WT. **h.** Schematic representation of the main calculated allosteric effect of F6.44M in the DA-D1-Gs complex (i.e. major path differences from D1WT). The legend describes the allosteric effects observed in the simulations.

Our analysis suggests that position 2.41 modulates the impact of N4.46 on signal transmission to the G-protein (**Fig.S28**). In D2, Y2.41 is engaged in strong polar contacts with N4.46 which locks its side-chain away from T2.37, preventing proper allosteric contacts with that residue and signal transduction (**Fig.4, Fig.S10**). In D1, F2.41 cannot establish such polar contacts but undergoes slight side-chain reorientation upon hydrophobic interactions with N4.46 (**Fig.S28**). This conformational shift enhances allosteric contacts with R55, N2.40, Y7.53 and F7.55, improving signal propagation down to the G-protein binding surface when compared to D1 WT (**Fig.S28**).

To further confirm that position 2.41 is a significant differentiator between D1 and D2, we characterized the D1 I4.46N-F2.41Y variant where F2.41 is mutated to the corresponding D2 residue. Consistent with our hypothesis, we observed a decrease in response to BRC and DA both in the context of WT and I4.46N D1 variants (**Fig. 5f, Fig.S26, Fig.S27**). Conversely, the D2 receptor incorporating the D1 residue at position 2.41 exhibited enhanced signaling activity (**Fig.S11**).

The DA-D1-Gs structure was also solved in complex with a Positive Allosteric Modulator (PAM) which considerably increases DA efficacy and binds near position 4.46^30^. Interestingly, our analysis revealed that the PAM and N4.46 have similar impacts on D1 structure, dynamics and allosteric signal propagation (**Fig.5g**, **Fig.S29**). These findings indicate that the intracellular local region involving TM2-TM4 may constitute a major determinant of the suboptimal DA efficacy for D1.

Investigation of D1 F6.44M revealed also critical differences with the same sequence variant in D2. While signal propagation through TM6 is blocked by M6.44 in D2, it continues along TM6 in D1, reaching M6.41 and the G-protein binding interface (**Fig. 5h**). Once again, the sequence-structure context of the receptor considerably determines the impact of M6.44 on ligand-mediated signal transductions. In M6.44’s neighborhood, the critical allosteric position 6.41 employs a Met in D1 instead of a Leu in D2. Due to its high conformational flexibility, the Met side-chain can accommodate the sequence variant at 6.44 and maintain optimal allosteric contact for signal transduction (**Fig.S30**). M6.44 also triggers long-range effects in the ligand binding site that strengthens allosteric communications between the ligand and G-protein binding interface, in agreement with the observed enhanced sensitivity for DA (**Fig.S30**). For instance, N6.55 undergoes a conformational change that shifts the binding contacts to DA and enhances the allosteric communication between the ligand and TM6 down to M6.44 (**Fig.S30**). A similar structural adaptation mechanism takes place on the other side of the binding pocket involving W7.43 which becomes a strong allosteric signal transmitter to M6.44 (**Fig.S30**).

To further validate our structural mechanisms and confirm that position 6.41 is a significant differentiator between D1 and D2, we characterized the D1 F6.44M-M6.41L variant where M6.41 is substituted to the corresponding D2 residue. Consistent with our hypothesis, we observed a decrease in ligand response when compared to D1 F6.44M (e.g. decreased DA efficacy, **Fig.S27**). Conversely, D2 receptors incorporating the D1 residue at position 6.41 exhibited enhanced signaling activity (e.g. increased BRC efficacy by close to 50%, **Fig.S12, Fig.S14**).

## Discussion

Overall, our study reveals that natural and synthetic ligands can regulate GPCRs by exploiting multiple intramolecular communication pathways emerging from distinct sets of residues at the receptor binding site. This multi-branched allosteric network architecture enables receptors to encode precise responses to chemically-diverse stimuli, in line with the considerable effects on ligand sensitivity and selectivity of even slight sequence alterations in these paths. Our data revealed a new alphabet of allosteric perturbations, showcasing the rich diversity of mechanisms by which receptors adapt to sequence variations through highly specific changes in both receptor and ligand structures and dynamics. Notably, ligand conformational flexibility and ability to bind to distinct allosteric sites within the binding pocket has emerged as a crucial and previously unforeseen mechanism for maintaining signal transduction through multiple allosteric pathways. This principle of ligand adaptation could be leveraged in the design of personalized medicines, where drugs might be tailored to accommodate patient-specific polymorphisms.

We also discovered that the receptor sequence-structure context has a significant impact on ligand potency and efficacy. These properties are under strong but distinct evolutionary pressures in dopamine D1 and D2 receptors which maintain suboptimal efficacy and potency, respectively.

Our study validates a computational framework for predicting allosteric perturbations and provides a mechanistic blueprint for inferring the impact of sequence polymorphism on receptor signaling functions. The method is general and should be applicable to map ligand binding sites, signal transduction pathways, enhance pharmacogenomic studies and aid drug discovery for a wide range of proteins. Lastly, our findings pave the way for the computational design of receptors with fully programmable ligand-sensing functions, which could be used for enhanced diagnostics, imaging, cell engineering, and applications in basic research and synthetic biology.

## Acknowledgments

We thank the Barth lab members for helpful discussions.

## Authors contributions

P.B. designed the study; M.H. developed AlloCraft; M.H. and A.S. performed the MD simulations; M.H. and D.K. carried out the computational design calculations; A.O., D.K. and M.A. performed the experimental validation of the designs under the supervision of T.W. and P.B.; all authors participated in the analysis, interpretation of the results and wrote the manuscript.

## Data and materials availability

the AlloCraft code and example to run the method is available on github: https://github.com/barth-lab/AlloCraft. The other data are available in the manuscript or the supplementary materials.

## Materials and Methods

### D2 ligand clustering

A Dopamine D2 receptor ligand list was obtained from the CHEMBL database (https://www.ebi.ac.uk/chembl/) from which agonists were extracted. The physicochemical properties of all agonists were summarized using the JOELib cheminformatics library. The resulting data were clustered using density-based clustering (DBSCAN)^31^ and finally dimensionality was reduced using principal component analysis (PCA).

### TRP channel cell-based assay

HEK-293 cells stably expressing the TrpC4β channel (generously provided by Dr. Michael X. Zhu) were maintained in DMEM supplemented with 10% fetal bovine serum (FBS) and 50μg/mL geneticin as a selective antibiotic and grown at 37°C and 5% CO2. FLIPR Membrane potential assays (Molecular Devices) were performed as previously described^1^. Briefly, the assay relies on the detection of a membrane-permeable fluorescent dye coupled to a non-permeable quencher. The non-selective cation TRP channel changes the membrane potential upon activation by Gαi, which enables the selective influx of the dye. 24 hours prior to the assay, 150,000 cells/well were seeded into clear-bottom 96-well plates and were reverse transfected with an optimized quantity of receptor DNA (present in the pcDNA3.1+ vector) and 0.5μL Lipofectamine 2000 per well. Prior to reading the transfected plates, the media was removed and the FLIPR dye was applied. The relevant drug was transferred into the plates during plate reading on a Flexstation3 multi-mode plate reader and changes in fluorescence (emission at 535nm, excitation at 565nm) were measured for a period of a maximum of 4 minutes. Controls were removed and maximum fluorescence values were reported as function of the logarithm of the drug concentration in GraphPad PRISM10.

### GαS BRET-EPAC cAMP assay

HEK-293T cells (source: ATCC) were maintained in DMEM supplemented with 10% fetal bovine serum (FBS) and grown at 37°C and 5% CO2. Upon agonist stimulation of the dopamine receptor D1, GDP-GTP exchange promotes GαS dissociation from the Gβ and Gγ subunits. GαS subsequently activates adenylyl cyclase, which increases the concentration of cAMP in the cell. The well-characterized BRET sensor CAMYEL^32^ (cAMP sensor using YFP-EPAC-RLuc) based on the exchange protein directly activated by cAMP (EPAC) changes conformation and BRET ratios decrease upon cAMP increase. 24 hours prior to the assay, 75’000 cells/well were seeded into white-bottom 96-well plates and reverse transfected with an optimized quantity of DNA to match WT receptor levels as well as an optimized quantity of the CAMYEL biosensor DNA using 0,5uL Lipofectamine 2000 per well. Right before reading, the media was removed and cells were washed once with HBSS and 40uL HBSS was added in each well. Coelenterazine h was added on top of the cells and incubated for 5min. After a first read, drugs were added in each well and change in light emission was recorded using a Mithras2 multimode plate reader. Controls were removed and changes in BRET ratios were plotted as function of the logarithm of the drug concentration in GraphPad PRISM10.

### Enzyme-linked Immunosorbent Assay (ELISA)

To ensure all receptor variants express at a level like the WT receptor control, ELISAs were performed against the 3xHA N-terminal tag present on each receptor variant. For each of the aforementioned cell-based assay, an accompanying ELISA plate was also reverse transfected in parallel using the same conditions as the main assay plate. On the day of the assay, the media was removed from the wells and the cells were fixed with a 4% paraformaldehyde (PFA) solution for 10 minutes. Fixation was followed by a 2% bovine serum albumin (BSA) solution incubation, anti-HA mouse primary antibody incubation, and an anti-mouse secondary HRP-linked antibody incubation. Each for a period of 1 hour with three PBS washes between each step. SuperSignal chemiluminescent substrate (Thermo Fisher) was added to each well and plates were incubated for 5 minutes before a luminescent reading on a Flexstation3 plate reader. Mock-transfected wells were used to determine and subtract the baseline signal from the remaining wells.

### Ligand docking

Ligand docking was performed to obtain ligand-bound complex structures not characterized experimentally.

Dopamine docking onto the solved dopamine D2 receptor active state structure (6VMS^29^) was carried out using the IPHoLD Rosetta protocol^33^. Only the receptor chain was used during all steps; heteroatoms and additional chains were removed. The overall protocol consists of sequential coarse-grained docking coupled with structural relaxation, decoy clustering, high resolution docking, and ligand clustering steps. During the first docking and relax step 10,000 decoys are generated, of which the lowest 10% scoring are used in the subsequent structure clustering step to diversify target receptor conformation. High resolution docking was performed on the cluster centers of the largest 6 clusters, again generating 10,000 decoys for each cluster center model and only using the lowest 10% scoring decoys for subsequent analysis. An additional filter was added to only keep the lowest 50% scoring decoys in terms of interface energy (interface_delta). The remaining decoys were used for ligand binding mode clustering using a

DBSCAN algorithm with ligand heavy atom coordinates as the input. The largest binding mode cluster was designated as the putative native binding mode and used for further analyses.

#### Ligand binding contacts from MD trajectories

Ligand contacts were calculated between non-Hydrogen atoms using a dual cutoff scheme. Two atoms get into contact when they are within 3.5 Å of each other and stay in contact until they are further than 5 Å apart. This scheme is used to remove high frequency noise from distance fluctuating around the distance cutoff.

#### Principle component analysis (PCA) of ligand poses from MD trajectories

PCA was performed on the cartesian coordinates of the non-hydrogen atoms of the ligands (dopamine, bromocriptine, and apomorphine) by combining trajectories from all from all the studied systems (**Fig.S18, Fig.S24, Fig.S25**) Representative models from the molecular dynamics trajectories were chosen as the highest density points in the space of principal components (PCs) 1 and 2.

### AlloCraft pipeline

#### 1. Prediction of allosteric properties

Prediction of allosteric properties is divided into two parts, structural and dynamic. AlloCraft analyzes dihedral information from molecular dynamics simulations and extracts:

- Mutual information (MI) followed by path clustering to get allosteric pathways (dynamic contribution to allostery)
- Kullback-Leibler divergences (KL) to represent perturbation response (structural contribution to allostery)

##### 1.1. Mutual information (MI) calculation

We calculated mutual information (MI) from correlated motions extracted from MD simulations (**see above**)^34,35^. To calculate MI from torsional angles, a list of all backbone (*φ* and *ψ*) and side chain (*Χ1* up to *Χ5* where applicable) torsion angles was built from the initial structure, the torsions were then extracted every 100 ps after removing the first 50 ns of every replica. The torsions were then histogrammed using 50 bins (50 x 50 bins for 2-dimensional histograms), and the marginal entropy was calculated using the following equation:

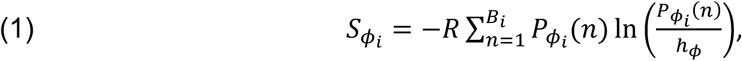

where 𝜙_*i*_ is the torsion being sampled for residue 𝑖, *R* is the gas constant, 𝐵*_i_* is the number of bins. 𝑃_𝜙_*_i_* (𝑛) is the probability of finding 𝜙*_i_* in bin 𝑛 defined as: 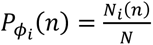, with *N_i_*(𝑛) being the number of datapoints/snapshots where 𝜙*_i_* falls in bin 𝑛 and *N* the total number of datapoints/snapshots. ℎ_&_ is the width of each bin in the histogram defined by the torsion 𝜙*_i_*. For two torsions 𝜙*_i_* and 𝜓*_j_* belonging to residues 𝑖 and 𝑗, the joint entropy is therefore defined as:

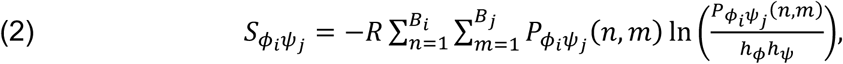

where 𝑃_𝜙*i*𝜓*j*_(𝑛, 𝑚) is the joint probability of finding 𝜙*_i_* in bin 𝑛 and 𝜓*_j_* in bin 𝑚. We then get the corresponding mutual information term 𝐼_𝜙*i*𝜓*j*_:

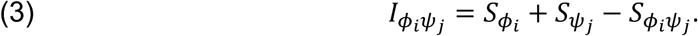

Correction for finite-size effects was also added:

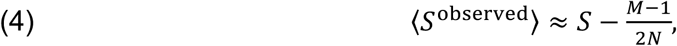

where 〈𝑆^observed^〉 is the estimated entropy using *N* datapoints and *M* is the number of histogram bins with non-zero probability^36,37^.

Another side effect of finite sample sizes is nonzero mutual information in independent datasets. To correct for this effect, we divide the observed MI space into 100 bins. In each MI bin, we randomly pick 5 dihedral pairs (or less if the bin has less than 5 samples) to represent the bin, and then for every dihedral pair 𝜙_*i*_and 𝜓_0_, we shuffle the time series of one of the observed dihedrals and recalculate MI with the shuffled dihedral. This process is repeated until the shuffled dihedral MI converges, and then the average of the resulting MI over the chosen dihedral pairs approximates the nonzero independent MI for a given MI bin. This value of independent MI is then subtracted from all MI values belonging to the bin. These permutations can also be used as a test of significance for MI, and the percentage of MI values from the various permutations that are larger than the observed MI approximates a *p*-value^38^. We used an MI significance level of *p* < 0.01 in our analysis.

Finally, entropies were tested for convergence by plotting 1^st^ and 2^nd^ order entropies as a function of number of frames. 1^st^ order entropy is defined as the sum of marginal entropies of all individual dihedrals (*φ, ψ*, and *Χ*’s) and 2^nd^ order entropy is the sum of joint entropy of dihedral pairs formed by the top 300 dihedrals with the highest summed MI.

##### 1.2. Allosteric pathway and pipeline calculation (see also Fig.S2)

Allosteric pathways were calculated by first constructing a graph where nodes are residues (from either peptide or receptor) and where edges are formed between any pair of residues with significant MI that have their Cαs within 10 Å. We then constructed pathways by minimizing the number of intermediate nodes while maximizing the sum of edge MIs between pairs of residues with significant MI whose Cαs further than 10 Å apart using Dijkstra’s shortest path algorithm^39^. Allosteric pathways were then sorted according to the MI of their terminal residues, and the number of pathways considered for further analysis in every system accounted for 85 % of cumulative mutual information. To cluster pathways into allosteric pipelines according to their closeness in the 3D structure, we define an overlap parameter, which is the percentage of nodes from two given pathways within a cutoff distance (7.5 Å). Overlapping allosteric pathways are clustered into allosteric communication pipelines using hierarchical clustering, and the strength of a pipeline is the MI weighted sum of pathways passing through it, where every pathway is weighted by the MI value of the pathway’s end residues. We considered the top-ranking 10 pipelines for our analysis. Allosteric residues or hubs are defined as the residues with the largest number of MI-weighted allosteric pathways passing through them. These key residues, referred to as allosteric hubs, are formally defined as the residues with the largest number of MI-weighted allosteric pathways passing through them. Allosteric hubs are usually characterized by allosteric strength scores (or hubscores) ≥ 100.

Delta MI values in the main text figures are reported for the top 3 pipelines that are the main contributors to the dynamic allosteric communications between the ligand and G-protein binding surfaces.

##### 1.3. Perturbation response using Kullback-Leibler divergence (KL)

We calculated Kullback-Leibler divergence (KL) from distributions of dihedrals extracted from two sets of MD simulations, a reference ensemble and a target ensemble as explained in McClendon et al.^28^. A list of all backbone (*φ* and *ψ*) and side chain (*Χ1* up to *Χ5* where applicable) torsion angles was built from the initial structures, excluding torsions that exist in one set of simulations but not the other (due to an amino acid substitution, for example). The torsions were then extracted every 100 ps after removing the first 50 ns of every replica. The torsions were then histogrammed using 50 bins, and the local KL was calculated using the following equation:

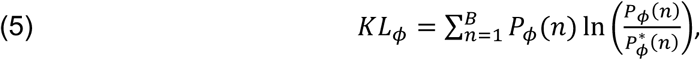

Where 𝜙 is the torsion being sampled, 𝐵 is the number of bins, 𝑃_𝜙_(𝑛) is the probability of finding 𝜙 in bin 𝑛 in the target ensemble, defined as mentioned in the MI calculation section, and 𝑃_𝜙_^∗^ (𝑛) is the probability of finding 𝜙 in bin 𝑛 in the reference ensemble.

The local KL for a single residue is simply the sum of the KL between reference and target ensembles for each of the applicable torsions for a given residue (𝜑, 𝜓, and 𝜒𝑠):

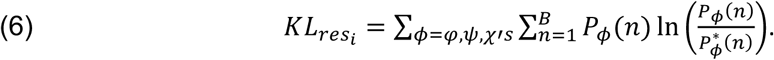

Despite the similarity of this expression to that of the marginal entropy (𝑆_𝜙δ_) mentioned before, it is a distinct thermodynamic quantity that measures the dissimilarity of two probability density functions (pdfs), while the marginal entropy 𝑆_𝜙*i*_ measures the level of disorder of a particular pdf. This makes local KL a suitable criterion for quantifying degree of freedom specific responses to a perturbation to a reference system. The perturbations we deal with in this study are amino acid substitutions, ensembles with different ligand, and ensembles in different conformational states (active vs inactive).

Corrections to the local KL were applied to counteract the effects of sample variability due to finite sampling^28^. To do that, we calculate KL from the reference ensemble using a statistical bootstrapping approach, which we use for significance testing and for correcting the calculated KL values (similar to mutual information expansion explained before). Reference sample KL is calculated by dividing the reference simulation into 𝑛_D_ blocks, and then using half the blocks as a proxy “reference” ensemble and the other half as a proxy “test” ensemble. Any non-zero KL between these proxy sets will approximate the bias to KL coming from intra-ensemble variability. The average bias thus becomes:

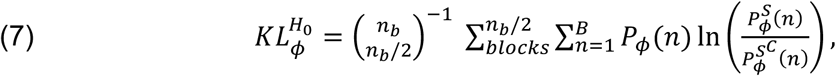

where 𝑆 are the subsamples constructed from half the blocks and 𝑆^J^ are the complementary subsamples from the other half of the blocks. We employ the distribution of proxy KL values to derive a *p*-value for the null hypothesis that the mean KL does not exceed what one would expect based on the variability observed in the reference ensemble. We then define a significance level 𝛼, which we use to zero out non-significant KL divergences, and subtract “excess” divergences from the significant ones. In this work, we set 𝑛_D_ = 6 and 𝛼 = 0.05.

#### 2. Computational design

Dopamine D2 variants were designed using RosettaMembrane^40^, a Rosetta-based protein structure prediction software utilizing a Monte Carlo gradient descent energy minimization algorithm enhanced with an anisotropic implicit membrane scoring function. The active-state dopamine D2 receptor structure (6VMS^29^) and the inactive state D2 receptor structure (6CM4^41^) served as the initial starting templates for the calculations. All heteroatoms and non-receptor or G-protein chains were removed from the starting structure. *In silico* scanning mutagenesis was carried out as follows:

1. Residues of interest were chosen using one of three criteria:

a. They belong to an allosteric pathway extracted from DA-bound and/or BRC-bound simulations. These positions are: L3.41, T5.54, V6.40, L6.41, F6.44, C6.47, V7.48
b. They maximize the difference in allosteric scores between ligands (BRC – DA).
c. positions are: for BRC biased hubs: I1.46, N1.50, V1.53, I4.46, V5.49, V5.53, L5.56, I5.59, A6.38, I6.39. For DA biased hubs: A2.54, A3.45, N7.45
d. For an expanded list of residues, we chose those that neighbor an allosteric hub or a conserved residue in class A GPCRs. We also scanned positions on TM4 and limited residues in loop regions. These positions are: V2.53, V2.66, E99 (ECL1), N143 (ICL2), V4.42, M4.45, S4.47, V4.49, W4.50, V4.51, L4.52, N176 (ECL2), Y5.58, V6.43, V7.44, A7.47, I7.51, I7.52, T7.54
2. Residues of interest were mutated and adjacent residues within 5Å were subjected to alternating cycles of sidechain repacking and backbone relaxation through Rosetta’s Monte Carlo-based energy minimization algorithm. 200 decoys were generated per design for each state to ensure score convergence. The lowest scoring decoys in inactive and active states were used for all subsequent analyses.
3. Perturbations in allosteric coupling from WT were estimated across the receptor structures using normal mode analysis (see below).

Designed mutations were selected for subsequent validation if:

a) the changes in inactive and active state structural energies were lower than 3 Rosetta Energy Units (REU) (a standard threshold used to filter out designs predicted to have significant energetic clashes or destabilizing effects):

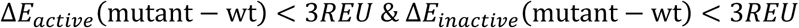

b) the mutation triggered significant changes in dynamic coupling in one versus the other state: ΔΔ ∑_*i*_ ∑_0_ 𝐷𝐶𝐶𝑀(𝑖, 𝑗), where 𝑖 and 𝑗 are allosteric residues, and the double delta is performed between (active and inactive) and (mutant and WT). This quantity has been related to the ligand-effector structural coupling ΔΔ𝐺^I^ under the assumption that ligand and receptor contributions to the energy between WT and mutant are similar^7^.

#### 3. Allosteric pathway coupling strength calculation

Global allosteric coupling through receptor structures was estimated as previously described^7^ using normal mode analysis. A “coupling” value for each receptor structure is derived from the dynamic couplings calculated for pairs of allosteric residues. These residues correspond to the major hubs that constitute the allosteric pipelines predicted by AlloCraft and through whose most of the clustered pathways transit (see above). Two lists of allosteric hubs were considered: 1) residues belonging to pathways common to both ligands; 2) Residues belonging to selective pathways engaged by only one of the ligands. In each case, dynamic cross-correlation matrices (DCCMs) are generated for the list of hubs using the lowest 20 normal modes which has been shown^42,43^ to correspond to large-scale, concerted allosteric motions. The sum total of the dynamic cross-correlations between pairs of allosteric hubs serve as the coupling value.

### Calculation of prediction success rate

The prediction success rate qualitatively reports on whether the method can predict gain or loss of function mutational effects. It was calculated using a combination of allosteric coupling strengths based on Normal mode analysis (NMA) of structures (i.e. DCCM values) and based on dynamic information extracted from MD simulations (i.e. MI values). If upon a mutation, the sign of the allosteric coupling strength matches either the change of efficacy or potency for both DA and BRC, then the prediction counts as a success. For the mutants that were not well predicted by allosteric coupling strengths from NMA but were simulated with MD, the difference of summed MI over all residues between mutant and WT was used to assess the success of the prediction.

### Molecular dynamics simulations

The starting structures used for MD simulations for dopamine D2 are: 1- BRC bound 6VMS^29^ with the last 20 residues of the C-terminal helix of Gi and the sequence re-mutated back to WT,

2- DA bound docked model based on 6VMS, and 3- AP bound model where the coordinates of AP were overlayed from AP-bound D1 structure (7JVQ)^44^ after alignment of TM region followed by relaxation with MD. For dopamine D1: 1- DA and 2- DA+PAM simulations used the DA structure 7CKZ^30^, while 3- BRC was overlayed from BRC-bound D2 structure (6VMS) after alignment of TM region followed by relaxation with MD. Β1AR structures are summarized in **Fig.S3d**. Missing loops were build using RosettaRemodel^45^. Mutant variants were generated based on the WT structures using RosettaMembrane^40^. The receptor-ligand-helix complex was inserted in a 90 × 90 Å^2^ POPC lipid bilayer solvated by 22.5 Å layer of water above and below the bilayer with

0.15 M of *N*𝑎^N^ and 𝐶𝑙^<^ ions using CHARMM-GUI bilayer builder^46,47^. Parameters for the two ligands (dopamine and bromocriptine) were generated using CGenFF^48^. Production simulations were performed with GROMACS 2019.4 with CHARMM36 forcefield^49^ in an *NPT* ensemble at 310K and 1 bar using a velocity rescaling thermostat (with a relaxation time of 0.1 𝑝𝑠) and Parrinello-Rahman barostat (with semi-isotropic coupling at a relaxation time of 5 𝑝𝑠) respectively. Equations of motion were integrated with a timestep of 2 𝑓𝑠 using a leap-frog algorithm. Each system was energy minimized using a steepest descent algorithm for 5000 steps, and then equilibrated with the atoms of the ligand-receptor-G-p helix complex and lipids restrained using a harmonic restraining force in 6 steps as shown in the table below:

| Step | 1 | 2 | 3 | 4 | 5 | 6 |
| --- | --- | --- | --- | --- | --- | --- |
| Time [ps] | 125 | 125 | 125 | 250 | 250 | 250 |
| Timestep [fs] | 1 | 1 | 1 | 2 | 2 | 2 |
| Ensemble | NVT | NVT | NPT | NPT | NPT | NPT |
| Pos restraints<br>[kJ/(mol·nm)] |  |  |  |  |  |  |
| Backbone | 4000 | 2000 | 1000 | 500 | 200 | 50 |
| Side chain | 2000 | 1000 | 500 | 200 | 50 | 0 |
| Lipid & ligand | 1000 | 400 | 400 | 200 | 40 | 0 |
| Dih restraints<br>[kJ/(mol·nm)] | 1000 | 400 | 200 | 200 | 100 | 0 |

After constrained equilibration, 5 to 10 replicas of 250 to 400 ns were run for each system. The first 50 ns of every simulation was discarded for equilibration of C-alpha RMSD, and the rest of the simulation was used for calculating statistics.

## Supplementary Figures

**Supplementary Figure 1.**
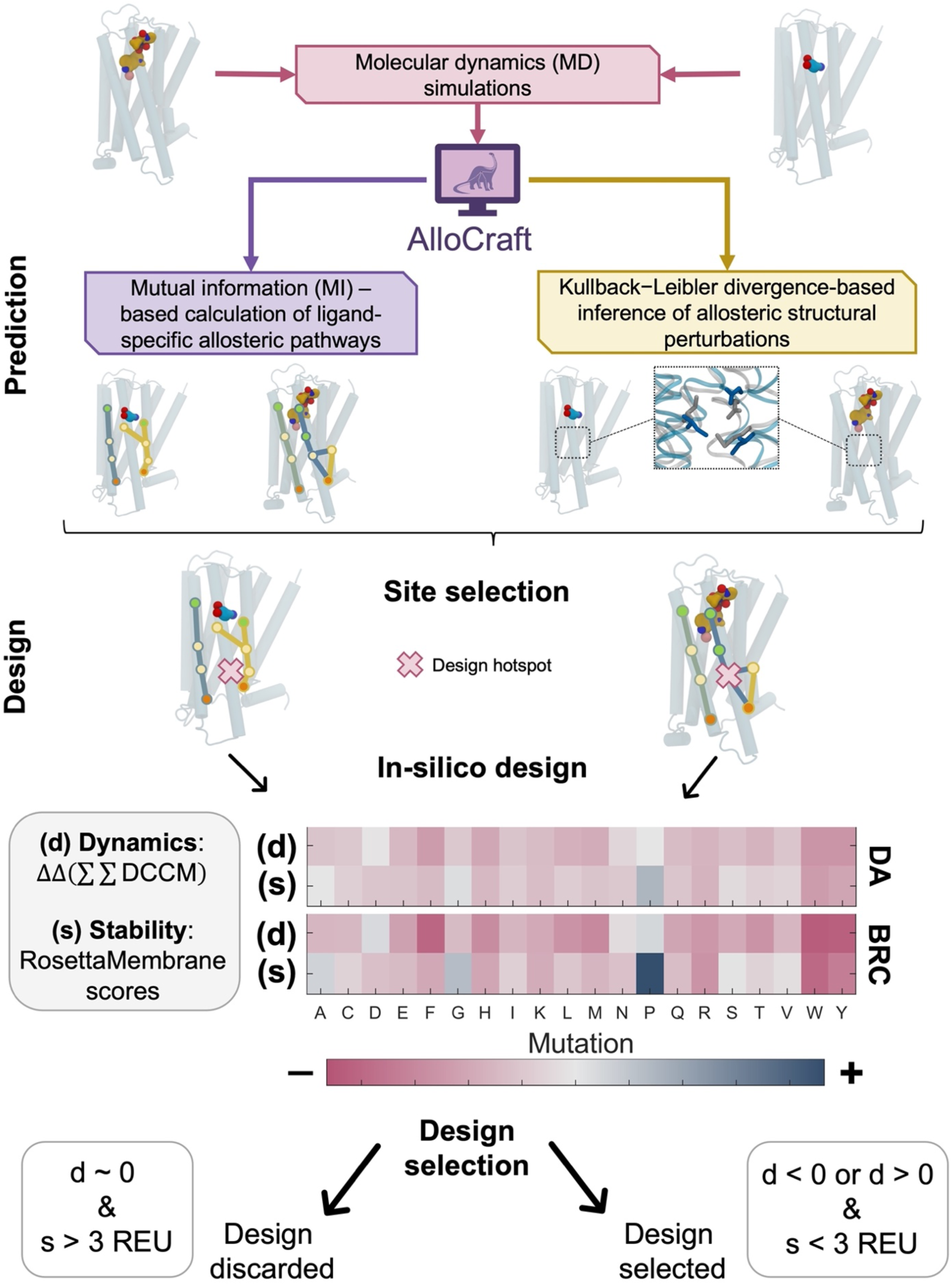
Schematic flowchart of AlloCraft. *<u>Prediction step.</u>* The first step of AlloCraft consists in predicting allosteric properties of GPCR-ligand systems. MD simulations are carried out for these complexes and peptide bond dihedrals are extracted from simulation trajectories. Dihedral information is processed to characterize either dynamic or structural allosteric properties (**Methods**). To infer dynamic allosteric coupling, AlloCraft calculates mutual information (MI) for each pair of residues. Residue networks (Pathways) are then constructed to connect highly coupled pairs that are more than 10 Å apart. The construction maximizes the MI along the path while minimizing its distance (i.e. number of residues connecting the distant pair). Constructed pathways are then clustered into allosteric pipelines (see **Fig.S2**). To infer allosteric structural perturbations, dihedral differences are calculated using Kullback-Liebler divergences between a reference and a target simulation. If the reference and target simulation correspond to an apo and ligand-bound states, respectively, then KLdiv will report the structural perturbations induced by ligand binding across the entire receptor. *<u>Design step.</u>* design hotspots are mutated to all 20 possible amino acids using RosettaMembrane. Amino acid substitutions are chosen based on stability (s) and correlated dynamics (d) scores (**Methods**).

**Supplementary Figure 2.**
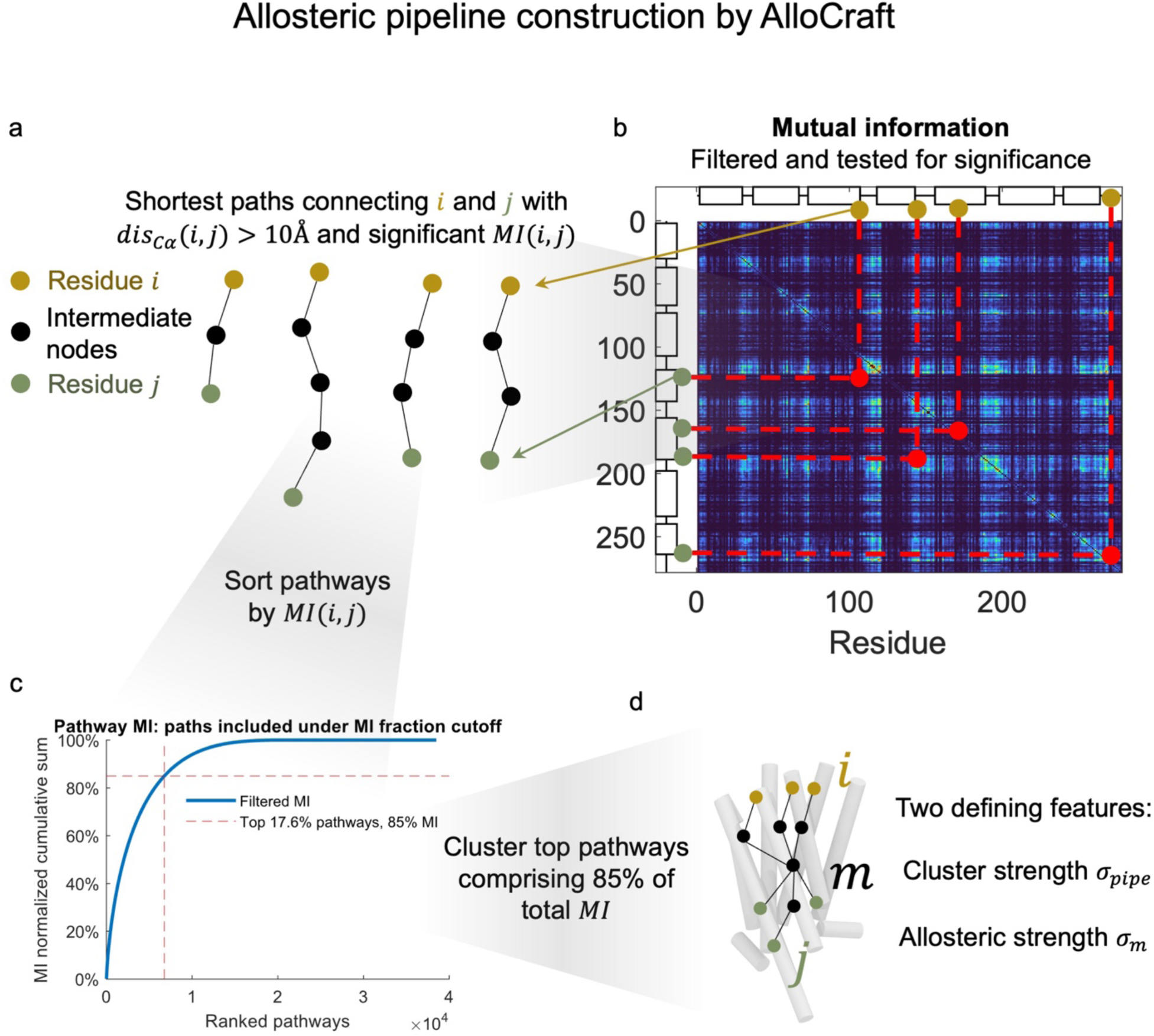
Construction of allosteric pipelines by AlloCraft from mutual information map (see Methods, AlloCraft pipeline). **a.** Residue pairs whose Cαs are further than 10Å apart and have significant MI between them are selected for pathway construction. Helices and loops are shown on the MI map as rectangles and lines respectively. **b.** Allosteric pathways are constructed by finding shortest paths that maximize edge MIs. **c.** Pathways are sorted by MI of their terminal residues. The top pathways that comprise 85% of cumulative MI are selected for clustering. **d.** An allosteric pipeline is a cluster of allosteric pathways with two defining features, the cluster strength, 𝜎_O.O?_, which equals the number of pathways belonging to this cluster, and this metric is used to rank allosteric pipelines. The second feature is for every residue *m* included in the cluster, and is the allosteric strength 𝜎_3_, which measures how many allosteric pathways contain residue *m* weighted by the pathway terminal 𝑀𝐼(𝑖, 𝑗).

**Supplementary Figure 3:**
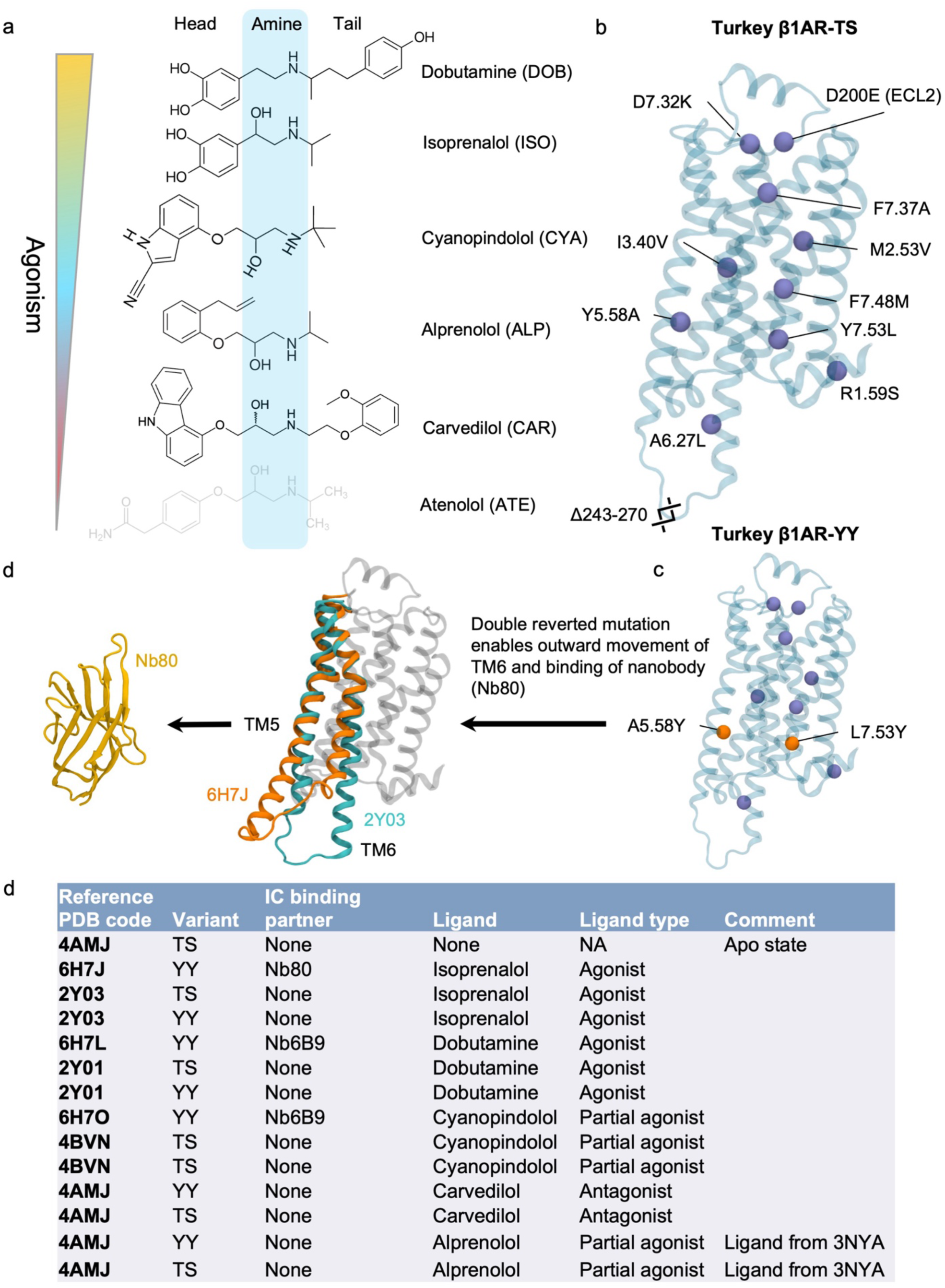
Ligand chemical structure and fitting of data to experimental efficacies of said ligands: **a.** Chemical structures of the *β*1AR ligands used in this study. The ligand atenolol was not simulated, but is present in the reference studies (Isogai et al., 2016 and Grahl et al., 2020). **b.** Turkey β1AR-thermostabilized variant (TS) showing mutations from WT, as reported in Isogai et al., 2016. **c.** Turkey β1AR-A5.58Y-L7.53Y (YY) reverted double mutant from β1AR-TS. **d.** Outward motion of TM6 enabled by the double reverted mutations, as evidenced by the slow tiemscale equilibrium in Grahl et al., 2020, which enables nanobody binding. **e.** Systems simulated for wild turkey *β*1AR. TS is the thermostabilized variant as described in Isogai et al. YY is the A5.58Y-L7.53Y reverted double mutant starting from TS. Ligands from different reference structures were overlayed after structural alignment and then relaxed with MD.

**Supplementary figure 4:**
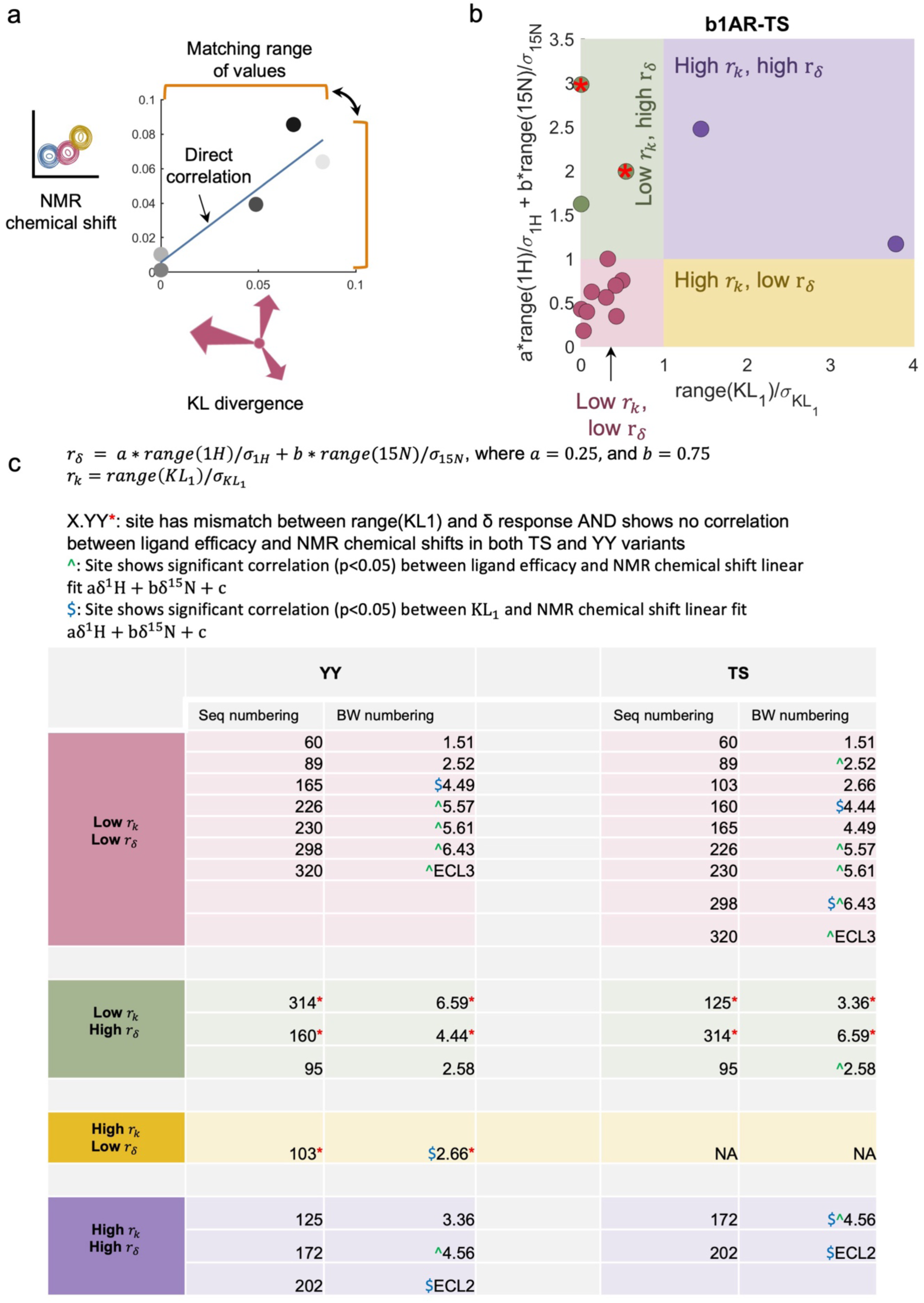
Qualitative comparison of KL-divergence and NMR chemicals shifts. **a.** Analysis of KL-divergence and NMR chemicals shifts by matching ranges of values (this figure) and direct correlation (shown in following figures) **b.** Scatter plot for the range of chemical shifts and range of KL-divergences for the thermostabilized (TS) mutant, σ represents standard deviation. **c.** Table showing the valine residues in beta1AR (wild turkey) that belong to the different classes described in Fig. 1. Legend of table in panel c: *r*_δ_ = 𝑎 ∗ *range*(1*H*)/𝜎_1H_ + 𝑏 ∗ *range*(15*N*)/𝜎_15*N*_ *r_k_* = *range*(*KL*_1_) X.YY*: site displays mismatch between range (KL1) and δ response AND shows no correlation between ligand efficacy and NMR chemical shifts in both TS and YY variants. ^: Site shows significant correlation (p<0.05) between ligand efficacy and NMR chemical shift linear fit aδ^1^H + bδ^15^N + c $: Site shows significant correlation (p<0.05) between KL_#_ and NMR chemical shift linear fit aδ^1^H + bδ^15^N + c

**Supplementary Figure 5:**
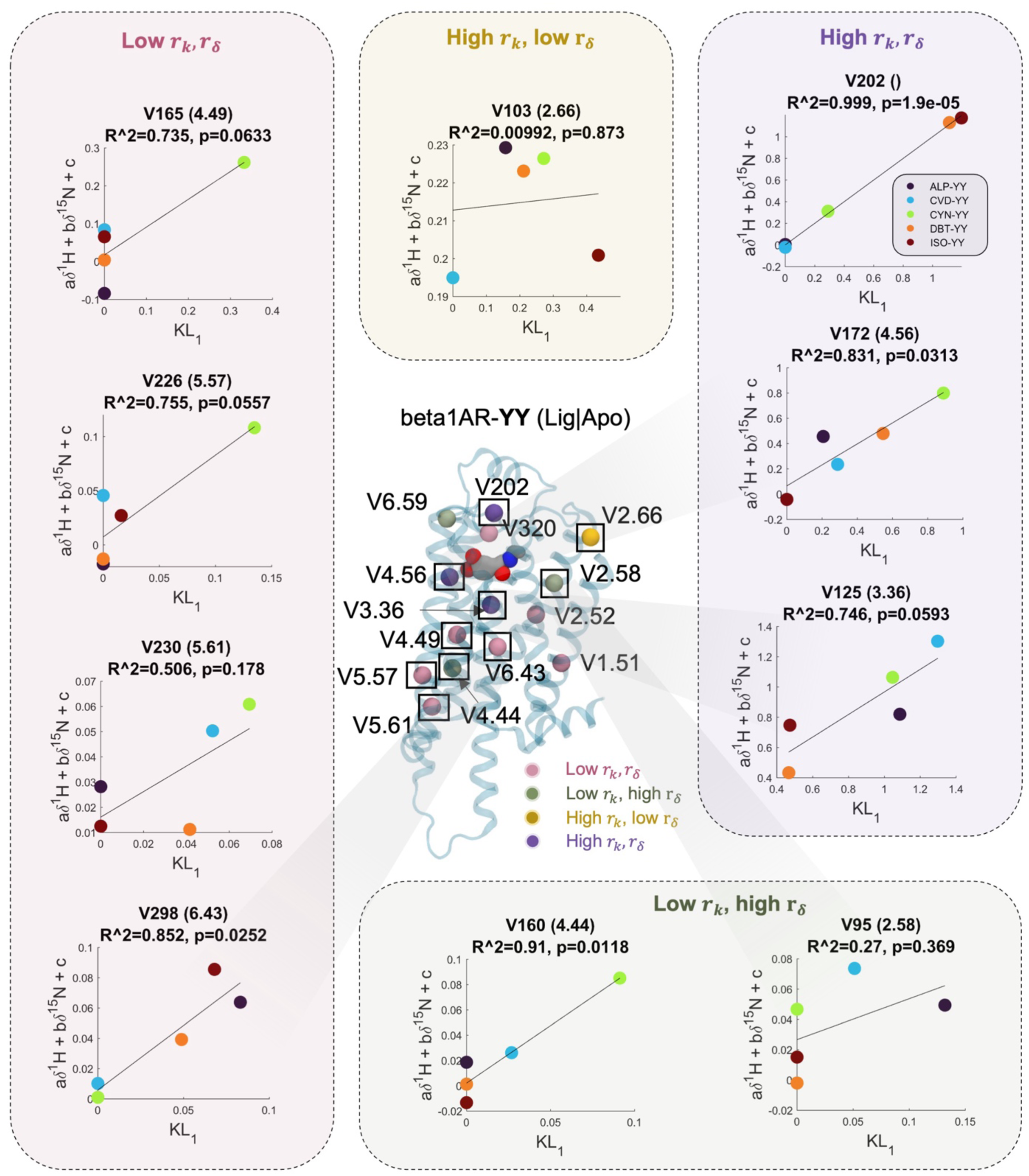
Correlation between KL-divergence and NMR chemicals shifts. Center. Valine residues in beta1AR (wild turkey) that belong to the different classes according to the range of chemical shifts and range of KL-divergences described in Fig. 1. b. and c. and Fig.S4. **Periphery.** Scatter plots for the correlation between KL-divergence and NMR chemical shift (as a linear combination of 1H, 15N, and a constant term) for the YY (A5.58Y/7.53Y) double reverted mutant. *r*_δ_ = 𝑎 ∗ *range*(1*H*)/𝜎_1H_ + 𝑏 ∗ *range*(15*N*)/𝜎_15*N*_, where 𝑎 = 0.25, and 𝑏 = 0.75, and *r*_k_ = *range*(*KL*_1_)/𝜎*_KL_*_1_

**Supplementary figure 6:**
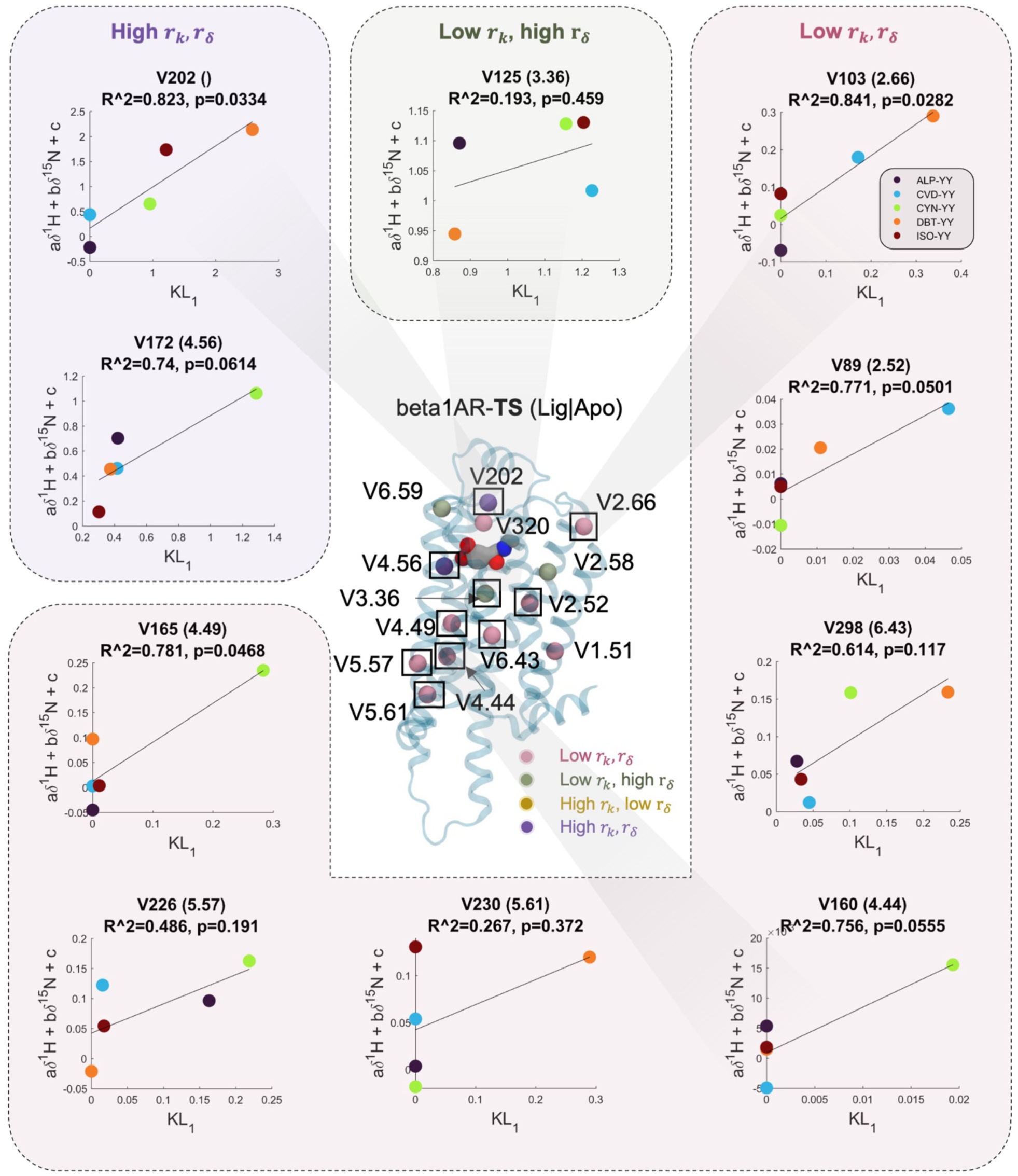
Correlation between KL-divergence and NMR chemicals shifts. Center. Valine residues in beta1AR (wild turkey) that belong to the different classes according to the range of chemical shifts and range of KL-divergences described in Fig. 1. **b.** and **c.** and Fig. S4. **Periphery.** Scatter plots for the correlation between KL-divergence and NMR chemical shift (as a linear combination of 1H, 15N, and a constant term) for the TS (thermostabilized) mutant.

**Supplementary Fig. 7.**
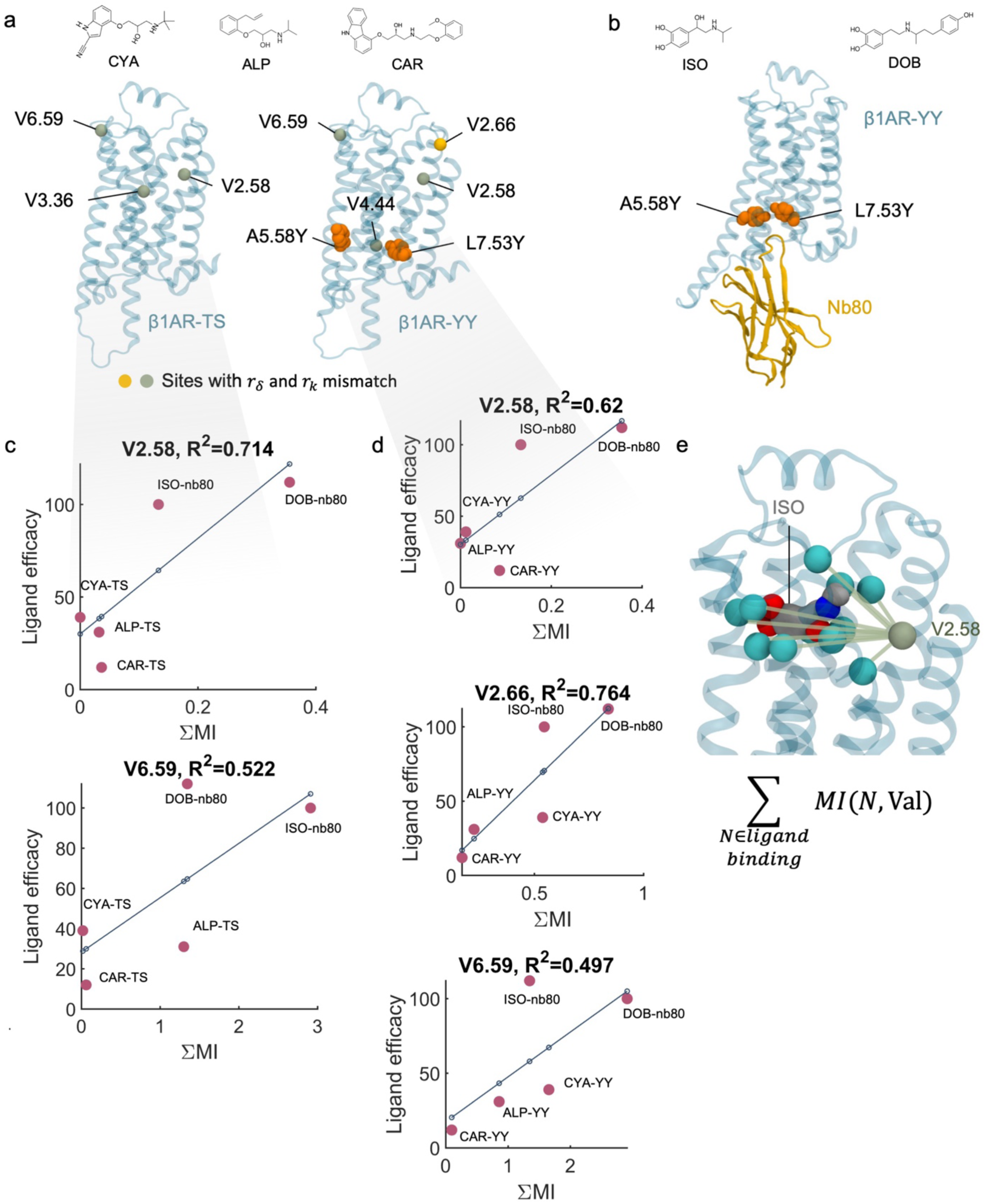
Fitting of mutual information to experimental efficacies of ligands for sites with. *r*_*δ*_ **and** *r_k_* **mismatch: a.** Sites with mismatched *r*_*δ*_ and *r_k_ for β*1AR-TS and *β*1AR-YY. Non-agonist ligands (CYA, ALP, and CAR, **Top**) were simulated as binary complexes with the receptor, while **b.** agonist ligands (ISO and DOB) were simulated as ternary complexes with Nb80 to simulate the receptor active state. Note that only the *β*1AR-YY construct can bind the nanobody and access the active state. **c-d.** G-s efficacy measured as percentage of activation of the full agonist isoprenalol plotted against summed MI between ligand binding residues and Valine residues in the B1AR TS (**c.**) and YY (**d.**) receptors. Valine sites (V3.36 for TS and V4.44 for YY) where MI did not display a signal (flat line) are not shown. **e.** Calculating mutual information sum between ligand binding residues (case of ISO shown) and target valine (case of V2.58 shown)

**Supplementary figure 8.**
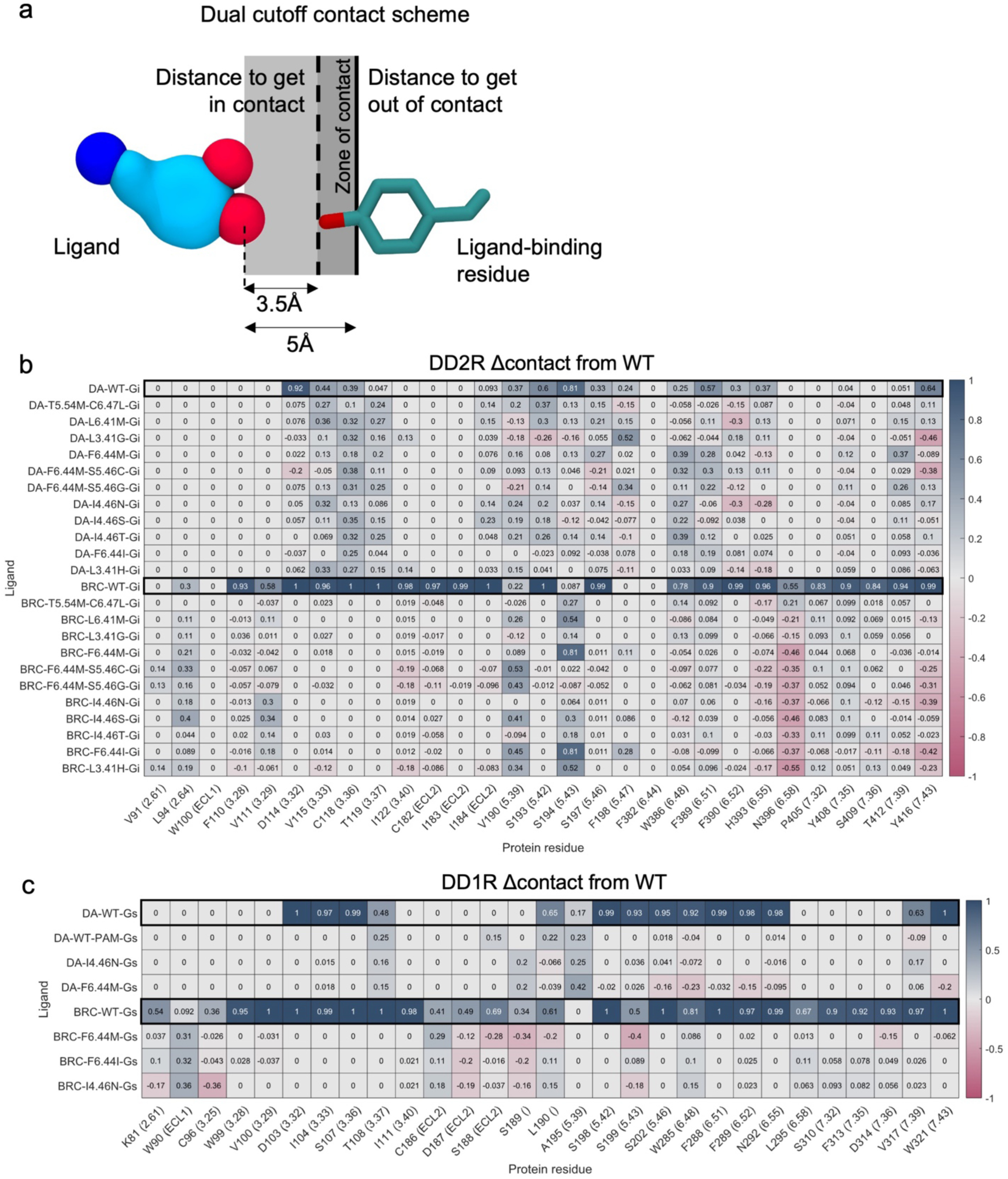
Ligand-receptor binding contacts analysis. **a.** Construction of contact map using a dual cutoff scheme: two heavy atoms (non-hydrogen) come in contact when they are within 3.5 A and leave contact when the atoms are further than 5 A apart. **b.** and **c.** WT values represent fraction of simulation in which the ligand is in contact with a given residue using the scheme defined in **a**. The values reported for the designs correspond to differences from WT. Panel **b.** contains data for DD2R, while panel **c.** for DD1R

**Supplementary figure 9.**
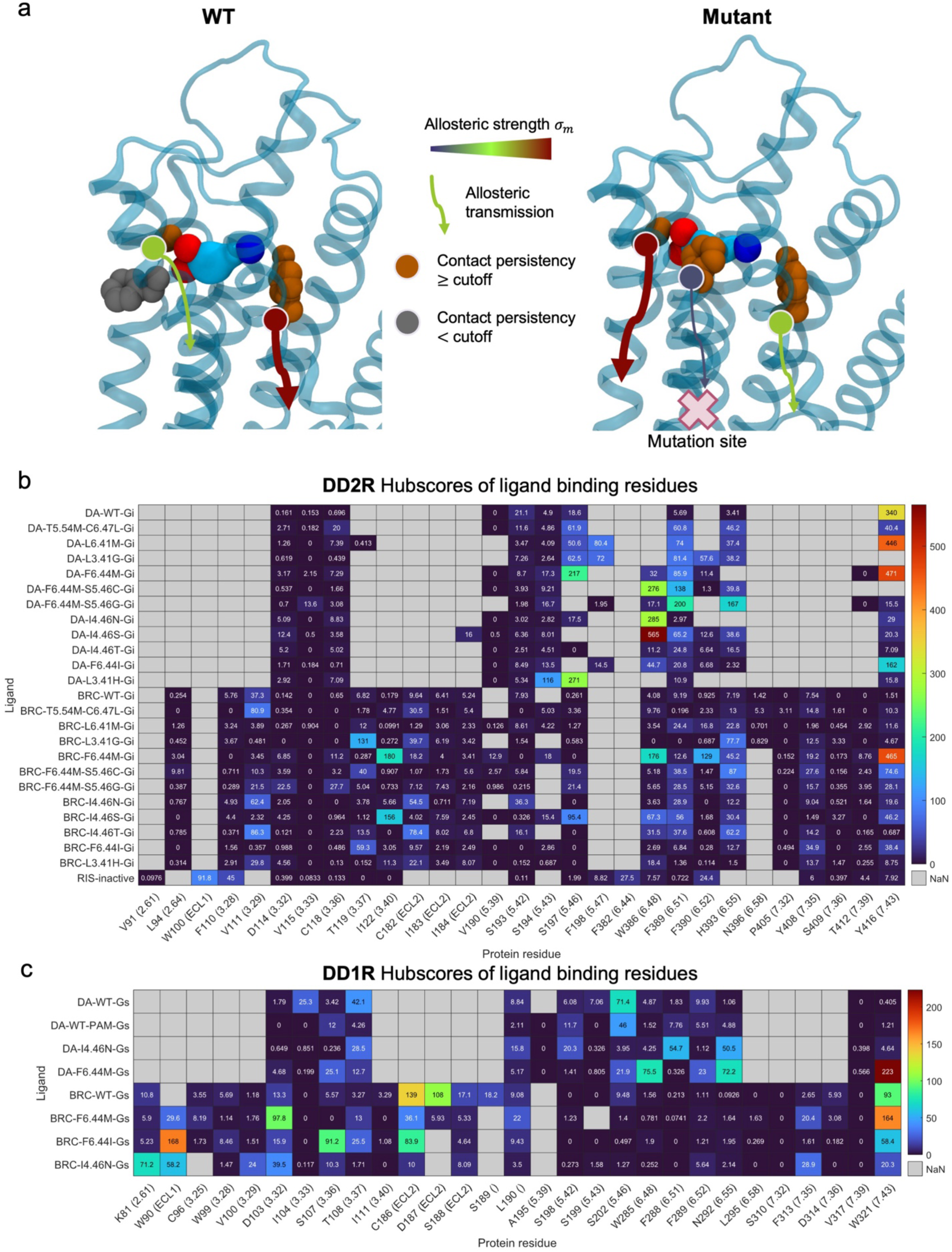
Ligand-receptor allosteric contacts analysis. **a.** Schematic representation of allosteric scores of ligand binding residues for WT (left) and a mutant (right). A mutation far from the ligand binding site can affect allosteric transmission from ligand binding residues. **b.** and **c.** Allosteric hubscores (see **methods**) of ligand binding residues for DA-bound, BRC-bound, and Ris-bound states. NaNs represent residues not having significant contact with the ligand. Significance cutoff is taken to be 30% of all simulated time. Panel **b.** contains data for DD2R, while panel **c.** for DD1R.

**Supplementary Figure 10.**
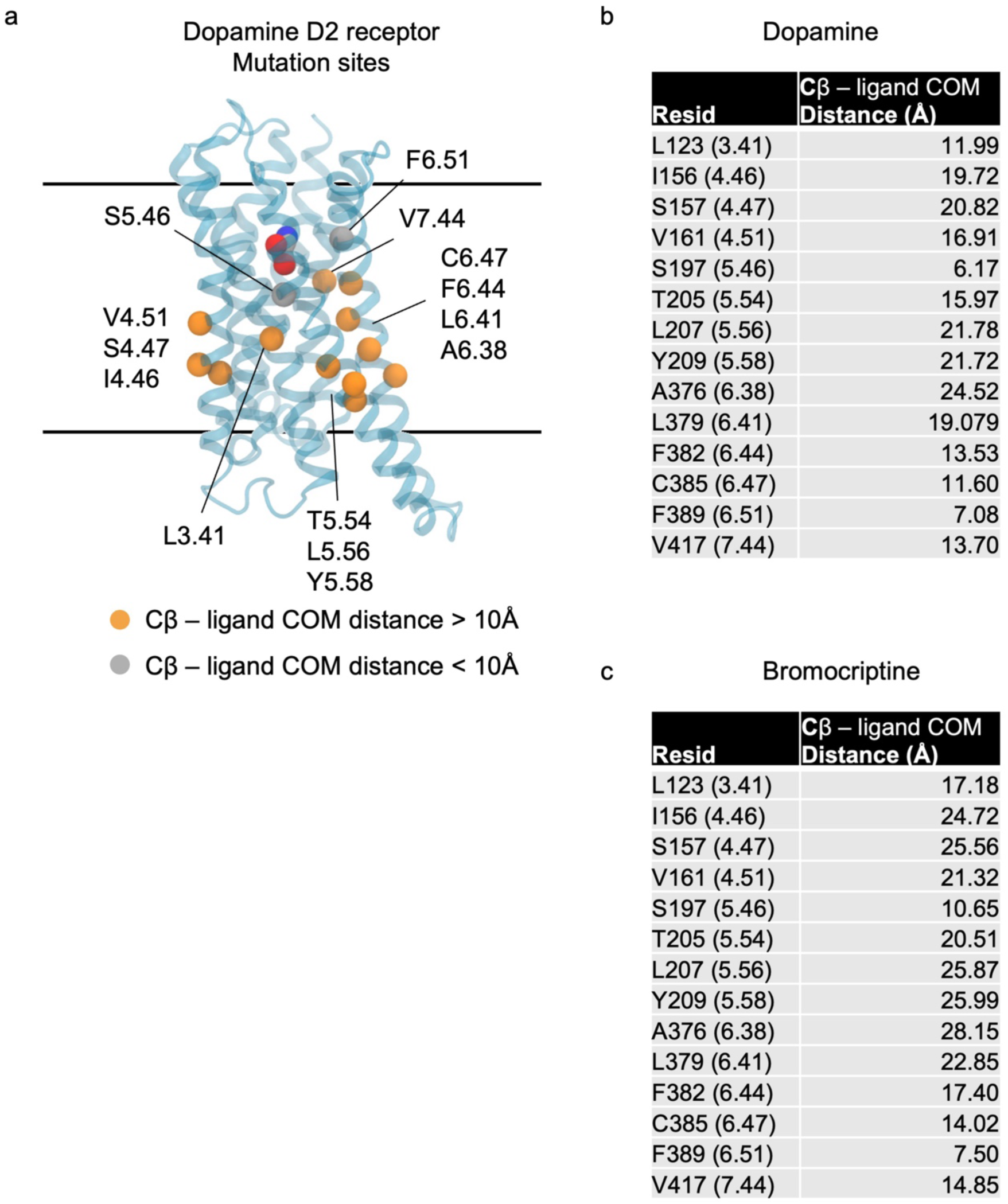
Position of D2 design sites. a. Positions of the mutated sites shown on the dopamine D2 receptor structure. b. Distances between Cβ of design sites and the center of mass (COM) of the ligand dopamine. c. Distances between Cβ of design sites and the center of mass (COM) of the ligand bromocriptine.

**Supplementary figure 11.**
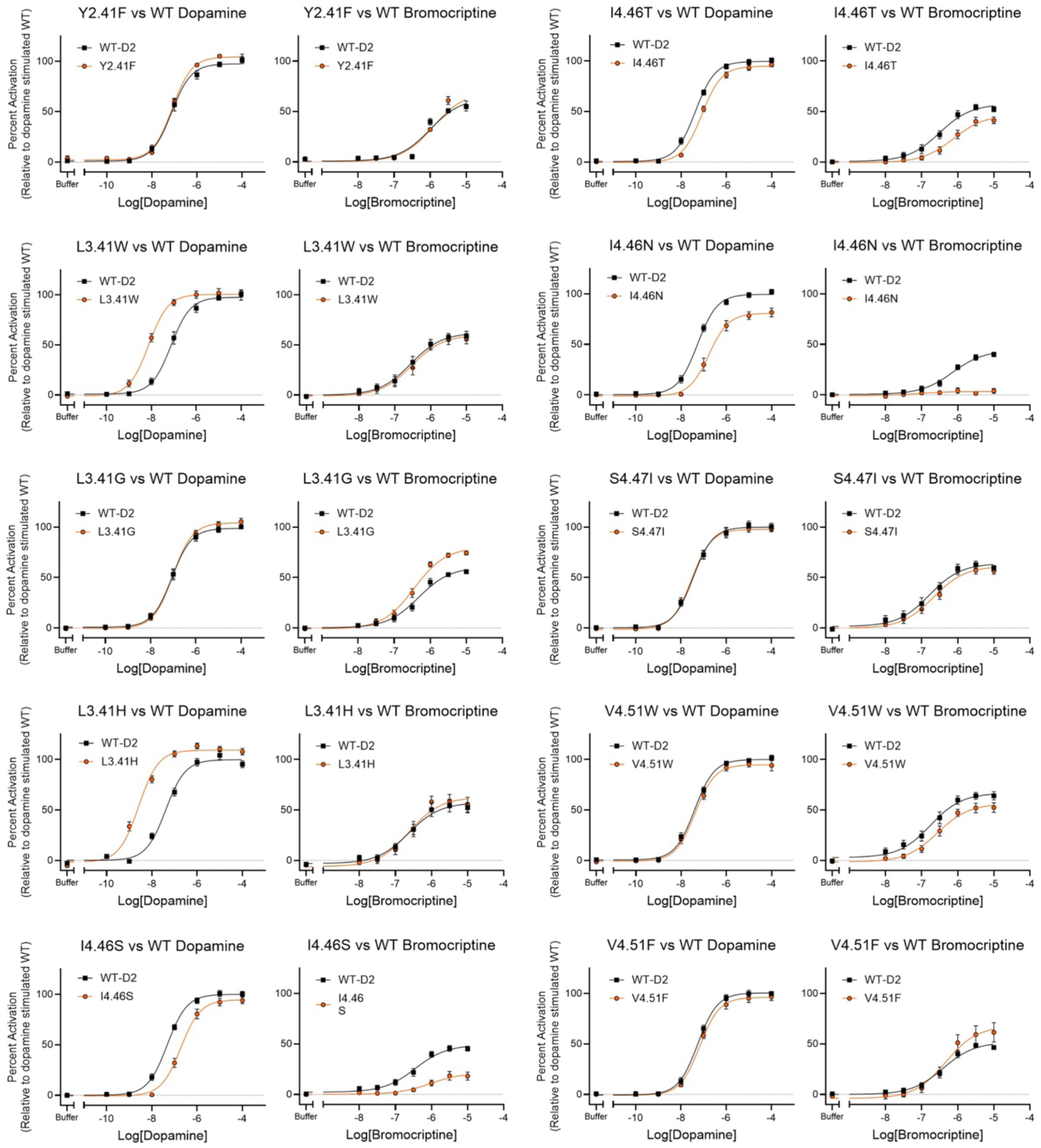
Experimental validation of the D2 designs. Ligand dose responses of D2 designed variants using the TRP channel assay assessing Gαi2 activation upon dopamine and bromocriptine binding to D2 receptors. WT D2DR in black and mutant D2DR in orange. Each point is the average of triplicate measurements from three independent experiments. SEM shown.

**Supplementary figure 12.**
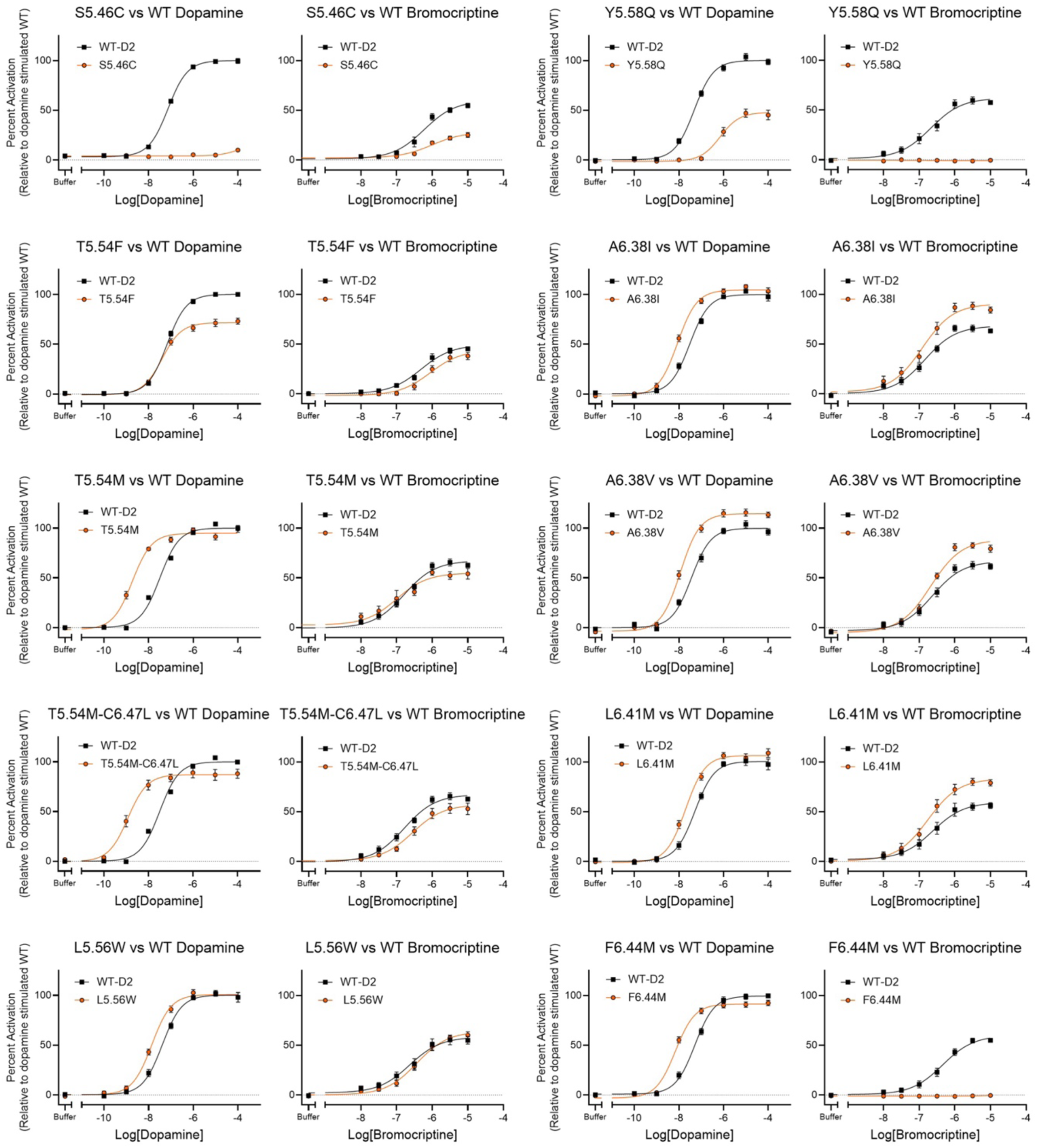
Experimental validation of the D2 designs. Ligand dose responses of D2 designed variants using the TRP channel assay assessing Gαi2 activation upon dopamine and bromocriptine binding to D2 receptors. WT D2DR in black and mutant D2DR in orange. Each point is the average of triplicate measurements from three independent experiments. SEM shown.

**Supplementary figure 13:**
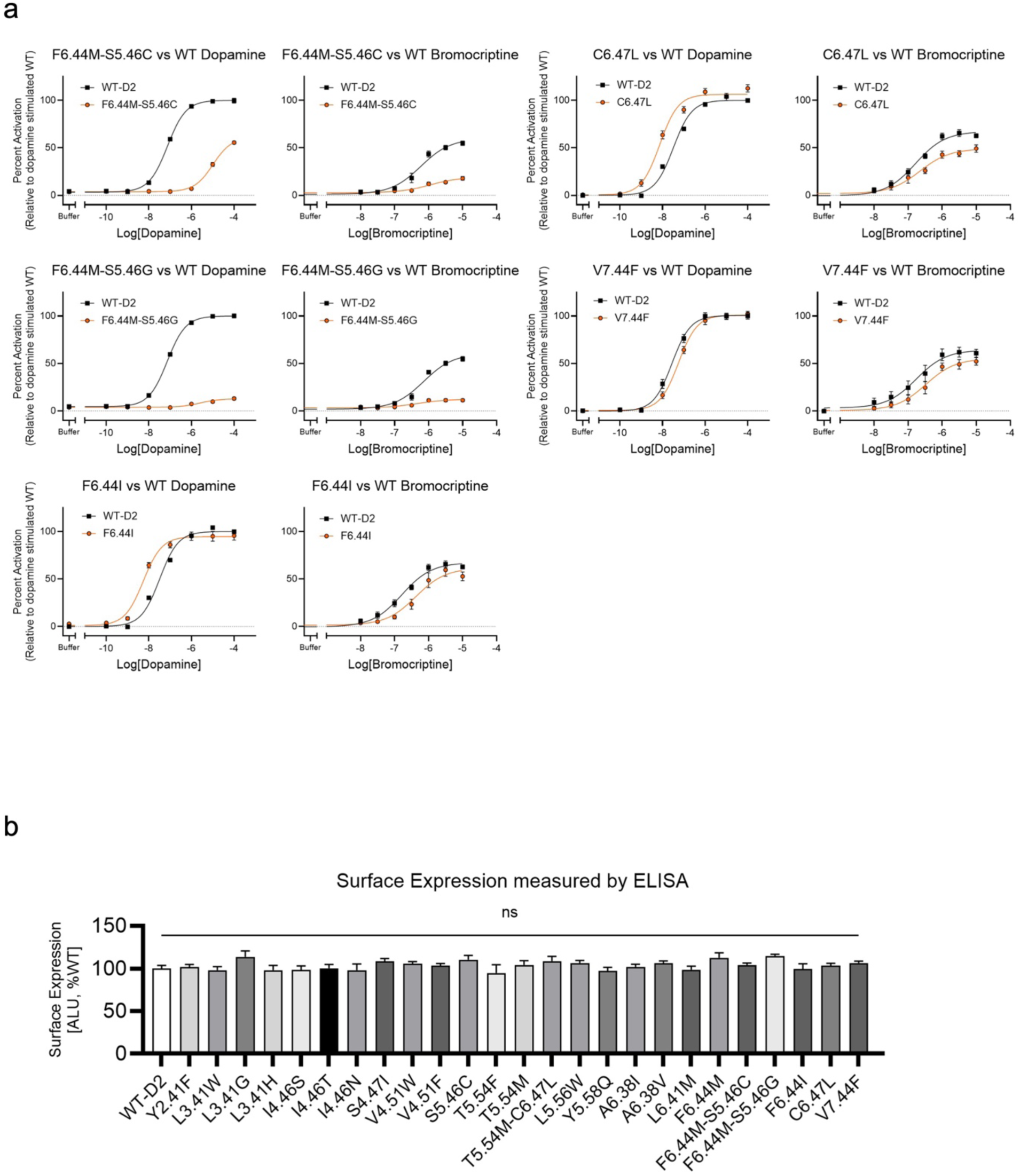
Experimental validation of the D2 designs. **a.** Ligand dose responses of D2 designed variants using the TRP channel assay assessing Gαi2 activation upon dopamine and bromocriptine binding to D2 receptors. WT D2DR in black and mutant D2DR in orange. Each point made in triplicate in three independant experiments. SEM shown. **b.** Surface expression of selected mutants of dopamine D2 receptor compared to WT-D2. Standard error of the mean shown. From 3 independent experiments, each made in triplicates.

**Supplementary figure 14.**
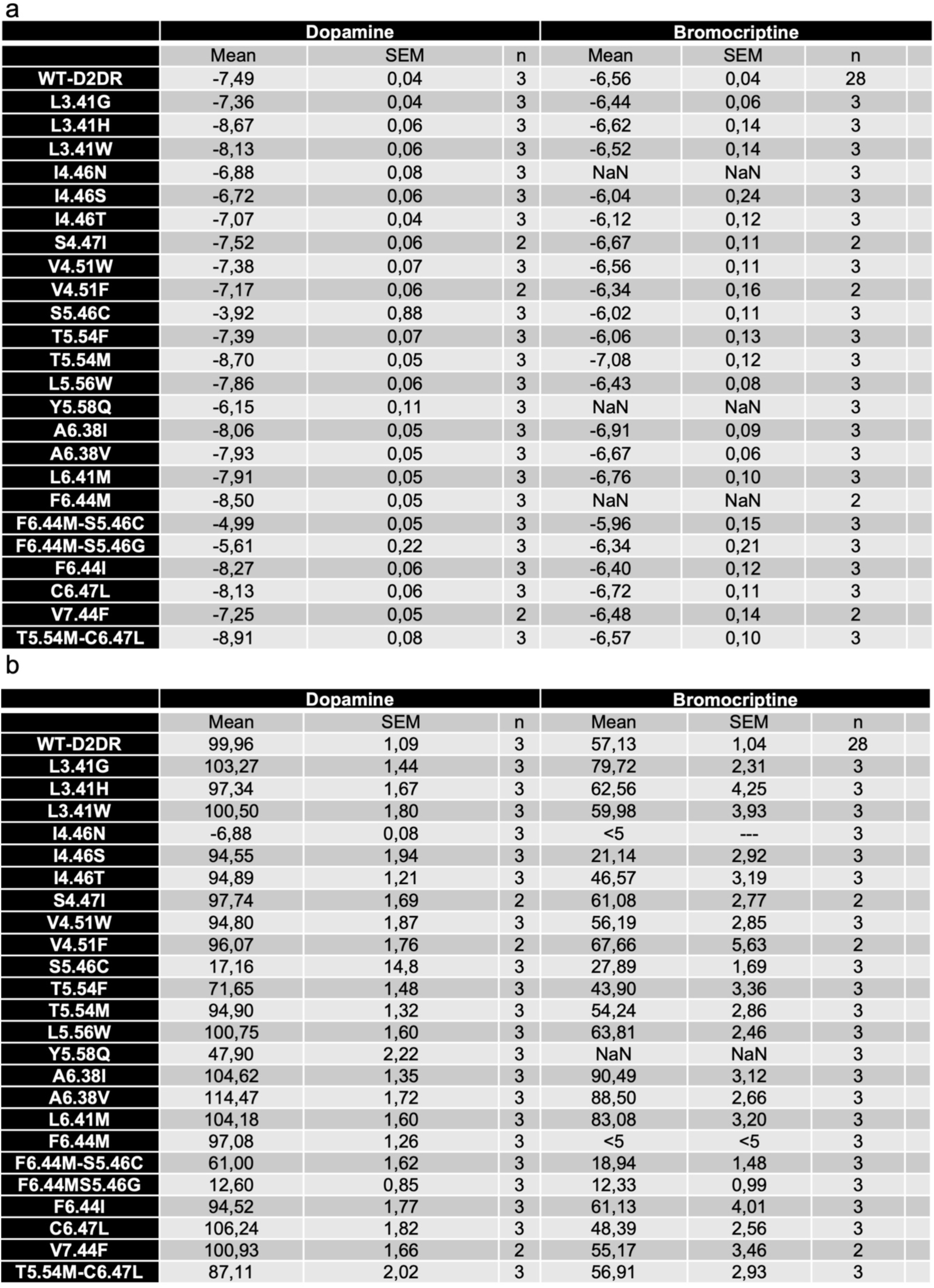
Ligand potencies and efficacies extracted from dose response curves. Dose response curves were fitted using GraphPad Prism software to obtain EC50 and efficacy. **a.** Mean log(EC_50_) and **b.** mean efficacies of selected mutants of dopamine D2 receptor. Standard error of the mean shown. n is the number of independent experiments, all made in triplicates. NaN: Experiment performed, no signal.

**Supplementary figure 15.**
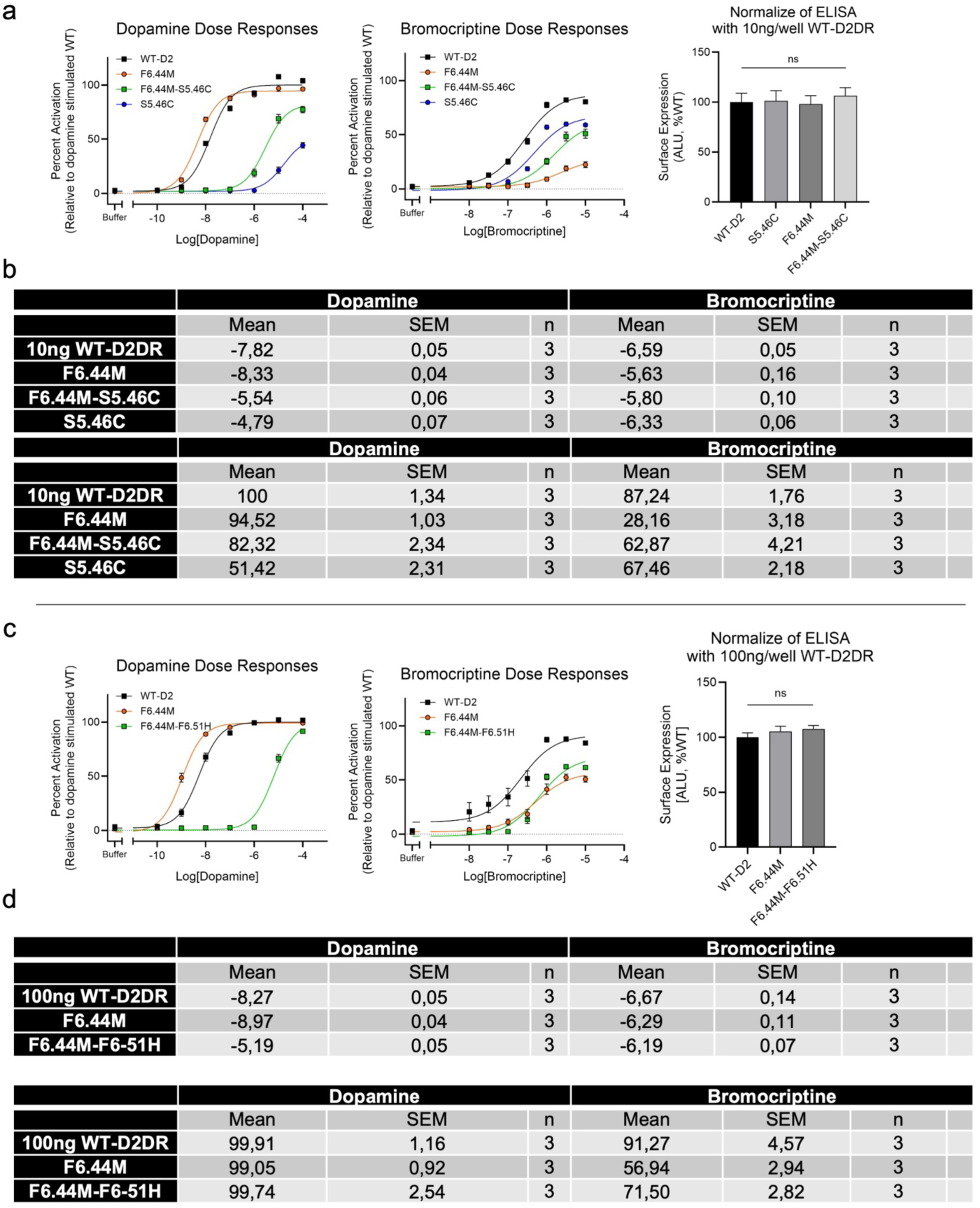
Experimental validation of D2 double mutant variants. a, **c.** Ligand dose responses of D2 designed variants using the TRP channel assay assessing Gαi2 activation upon dopamine and bromocriptine binding to D2 double mutant variants using increased amounts of transfected receptors and surface expression normalized to WT-D2. For the dose responses: WT D2DR in black and mutant D2DR in colours. Each point made in triplicate in three independant experiments. SEM shown. **b, d**. Mean log(EC50) and mean efficacies of selected mutants of dopamine D2 receptor extracted from the dose responses in a and c. Standard error of the mean shown. n is the number of independent experiments, all made in triplicates.

**Supplementary Figure 16:**
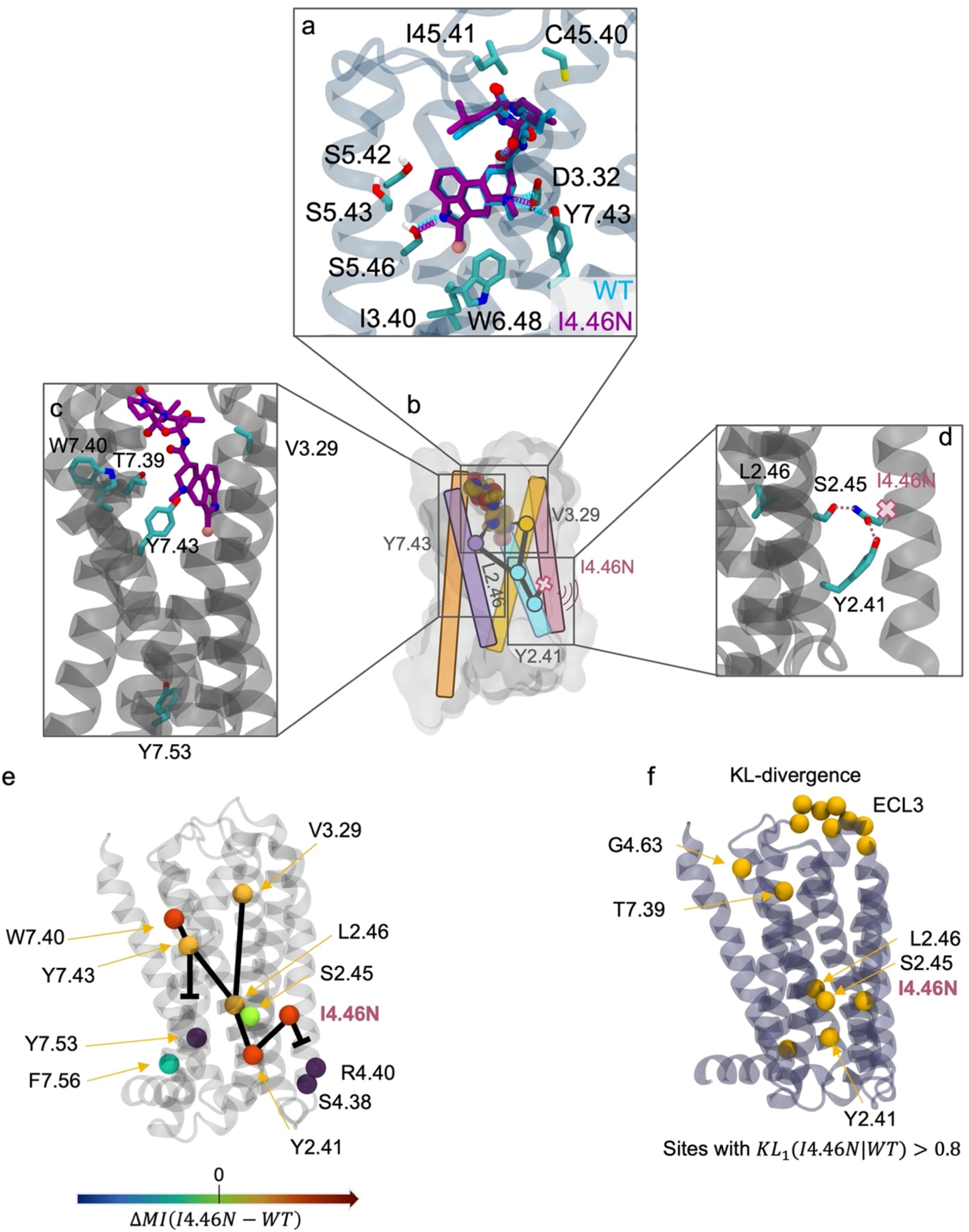
I4.46N impact on D2 allosteric response to BRC. **a.** Representative frame of highest density ligand binding pose for WT (cyan) and I4.46N (purple) extracted using PCA. **b.** Effect of the mutation is interpreted as an allosteric sink. **c.** zoomed-in view on ligand binding residues. **d.** zoomed-in view on the mutation site showing interaction network formed by the mutated residue. **e.** Sites with large differences in mutual information (MI) mapped on the DD2R structure. MI is calculated as a sum between a given residue and all other residues. The major path connection differences from WT are represented using straight black lines and broken paths using T lines. **f.** KL-divergence visualized on the structure for residues having KL_1_ > 𝟎. 𝟖. **Supplementary note for panel f.** Analysis of the calculated allosteric perturbations identifies the following major impacts of I4.46N on D2 allosteric responses to BRC: 1. N4.46 forms polar interactions with Y2.41 and S2.45, causing a change in their rotameric state distributions. 2. Consequently, Y2.41 becomes more dynamically coupled to L2.46 such that both residues act as allosteric hubs at the junction of allosteric pathways connecting the ligand binding region (V3.29, Y7.43) to TM2 down to N4.46. 3. Dynamic communication is therefore largely funneled through TM2 at the expense of TM4,5,6, and 7 which display reduced MI. 4. Despite the gain in dynamic communication along TM2 reaching TM4 at N4.46, this site becomes largely uncoupled to R4.40 and S4.38 and loses its ability to communicate with the intracellular surface. 5. Overall, we observe an important loss of dynamic communication (25% from WT) between the G-protein binding residues (located on helices 5, 6, and 7, and ICL2) and the rest of the receptor, consistent with the measured drop in G-protein activation. **Conclusion**: I4.46N acts as an allosteric sink, where the dynamic information is diverted toward the mutation neighbourhood away from the G-protein binding site.

**Supplementary Figure 17:**
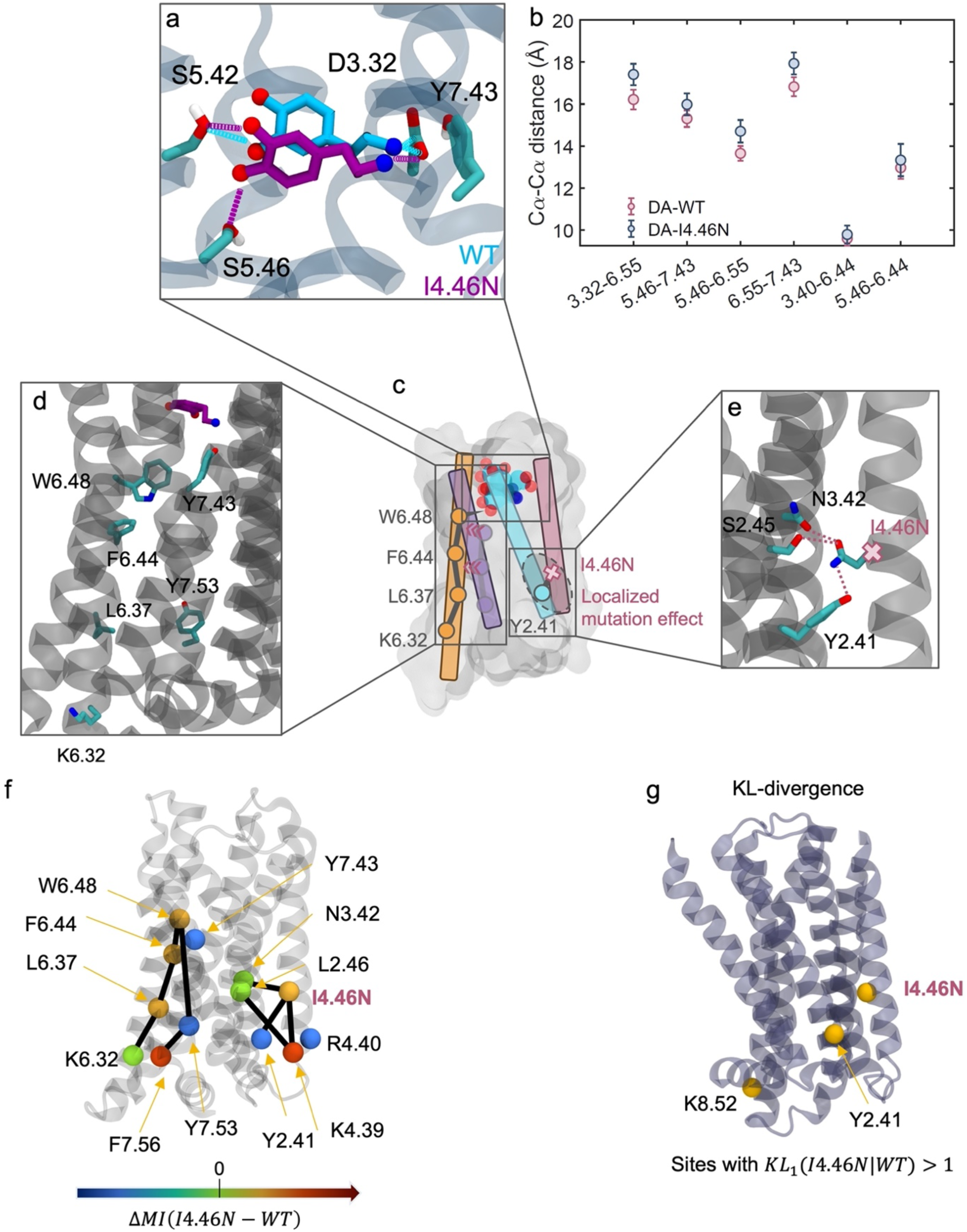
I4.46N impact on D2 allosteric response to DA. **a.** Representative frame of highest density ligand binding pose for WT (cyan) and I4.46N (purple) extracted using PCA. **b.** Distances for ligand binding residues and PIF motif layer for DA WT and I4.46N. Means and SEM are shown. Simulation frames were taken from the most populated DA binding cluster as identified by PCA (**methods**) **c.** The mutation acts as an allosteric sink and triggers a conformational repositioning of DA in the binding site (**a**) that is associated with a major path rewiring along TM6 reaching the G-protein binding surface. **d.** zoomed-in view on helices 6 and 7 showing residues involved in the allosterically rewired pathway. **e.** zoomed-in view on the mutation site showing interaction network formed by the mutated residue. **f.** Sites with large differences in mutual information (MI) mapped on the DD2R structure. The major path connection differences from WT are represented using straight black lines. MI is calculated as a sum between a given residue and all other residues. **g.** KL-divergence visualized on the structure for residues having KL_#_ > 1. **Supplementary note for panel f.** Analysis of the calculated allosteric perturbations identifies the following major impacts of I4.46N on D2 allosteric responses to DA: 1. N4.46 forms polar interactions with Y2.41 and S2.45, but it only causes a large change in their rotameric state distribution of Y2.41, as evidenced by KL-divergences. 2. However, unlike for the BRC-bound receptor, Y2.41 does not become an allosteric hub and loses coupling to its neighbors L2.46 and N4.46. N4.46 does not act as a sink but, instead, propagates dynamic information efficiently to the G-protein binding surface along TM4 through K4.39 to ICL2. 3. Importantly, N4.46 triggers significant long-range conformational changes in the extracellular binding site. DA accommodates these structural changes by shifting binding pose. 4. DA repositioning in the binding site changes a key ligand allosteric contact with the receptor from Y7.43 to W6.48. 5. Consequently, we observed a shift in allosteric communication from TM7 to TM6 as evidenced by pathway topology and summed MI per TM helix. **Conclusion:** I4.46N acts as a rewiring hub. It triggers significant conformational changes in the ligand binding site leading to a shift in DA binding pose and a rewiring of allosteric pathways from TM7 to TM6.

**Supplementary Figure 18.**
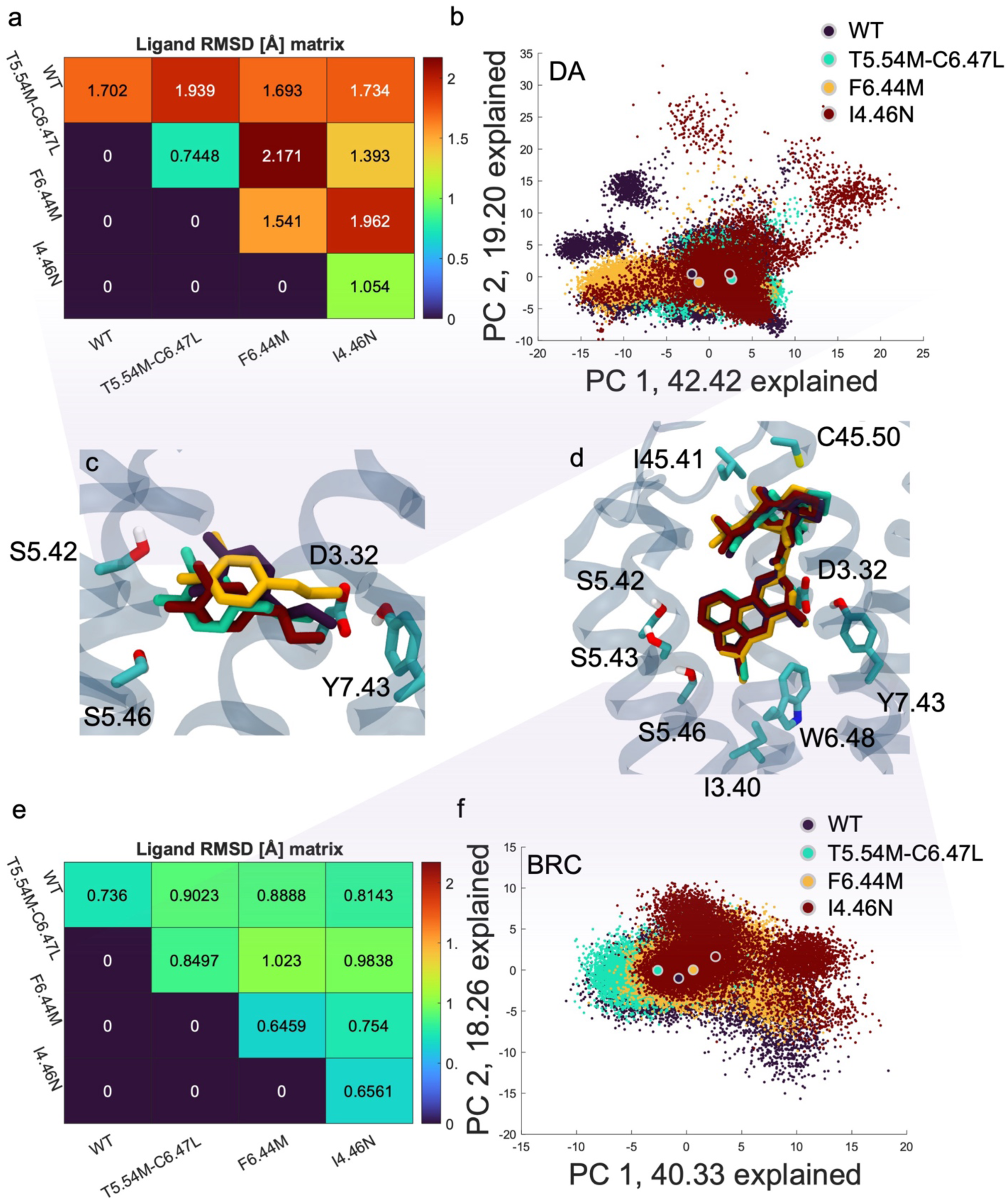
Conformational adaptation of ligands to D2 mutations. Principal component analysis of the ligand binding poses of dopamine (**a-c**) and bromocriptine (**d-f**) during molecular dynamics simulations for the following variants: WT, T5.54M-C6.47L, F6.44M, and I4.46N. The labelled dots represent the centroid of the PC spread for a given variant. The percentage of variance of the data explained by every PC is in the axis labels. RMSD is calculated by first finding highest density points for every system in PC space via histogramming (using 51 bins in every dimension). The 10 frames that are closest to the highest density PC point are then used to calculate RMSD. The mean is shown in the matrix

**Supplementary Figure 19:**
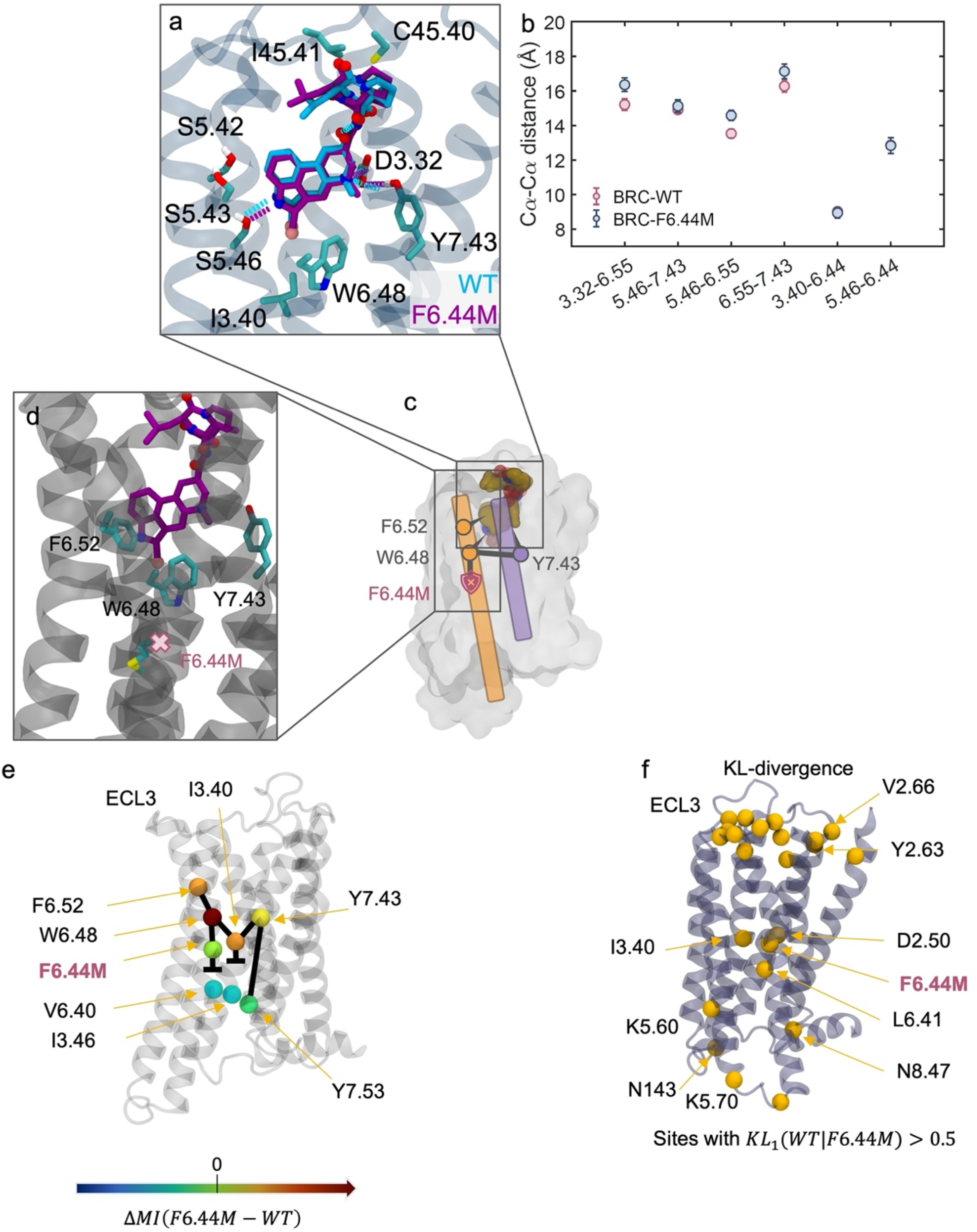
F6.44M impact on D2 allosteric response to BRC. **a.** Representative frame of highest density ligand binding pose for WT (cyan) and F6.44M (purple) extracted using PCA. **b.** Effect of mutation as an allosteric block **c.** zoomed-in view of helix 6 showing the residues transmitting the signal down to the allosteric block at the mutation site. **d.** Sites with large differences in mutual information (MI) mapped on the DD2R structure. MI is calculated as a sum between a given residue and all other residues. The major path connection differences from WT are represented using straight black lines and broken paths using T lines. **e.** KL-divergence visualized on the structure for residues having KL_#_ > 0.5. **Supplementary note for panel e.** Analysis of the calculated allosteric perturbations identifies the following major impacts of F6.44M on D2 allosteric responses to BRC: 1. We observed a complete loss of MI connections on TM6 around 6.44 and 6.48 and a significant decrease in allosteric communication through TM5. 2. 7.43 establishes new allosteric connections to 6.48 and 3.40. 3. Through the new 7.43-3.40 connection, 7.43 sends large amount of information to 6.48 which is not transmitted to the intracellular surface. 4. 7.43 connects to 7.49, which enables the restoration of WT-like communication through TM7. 5. Despite point #4, there is weaker allosteric coupling to 7.53 which is the hub distributing signal to the G-protein binding residues: 6.36 and 7.56. Conclusion: F6.44M triggers a complete loss of communication through TM6, weaker communication through TM5 while communication through TM7 is maintained to a certain level through local rewiring around 7.43.

**Supplementary Figure 20:**
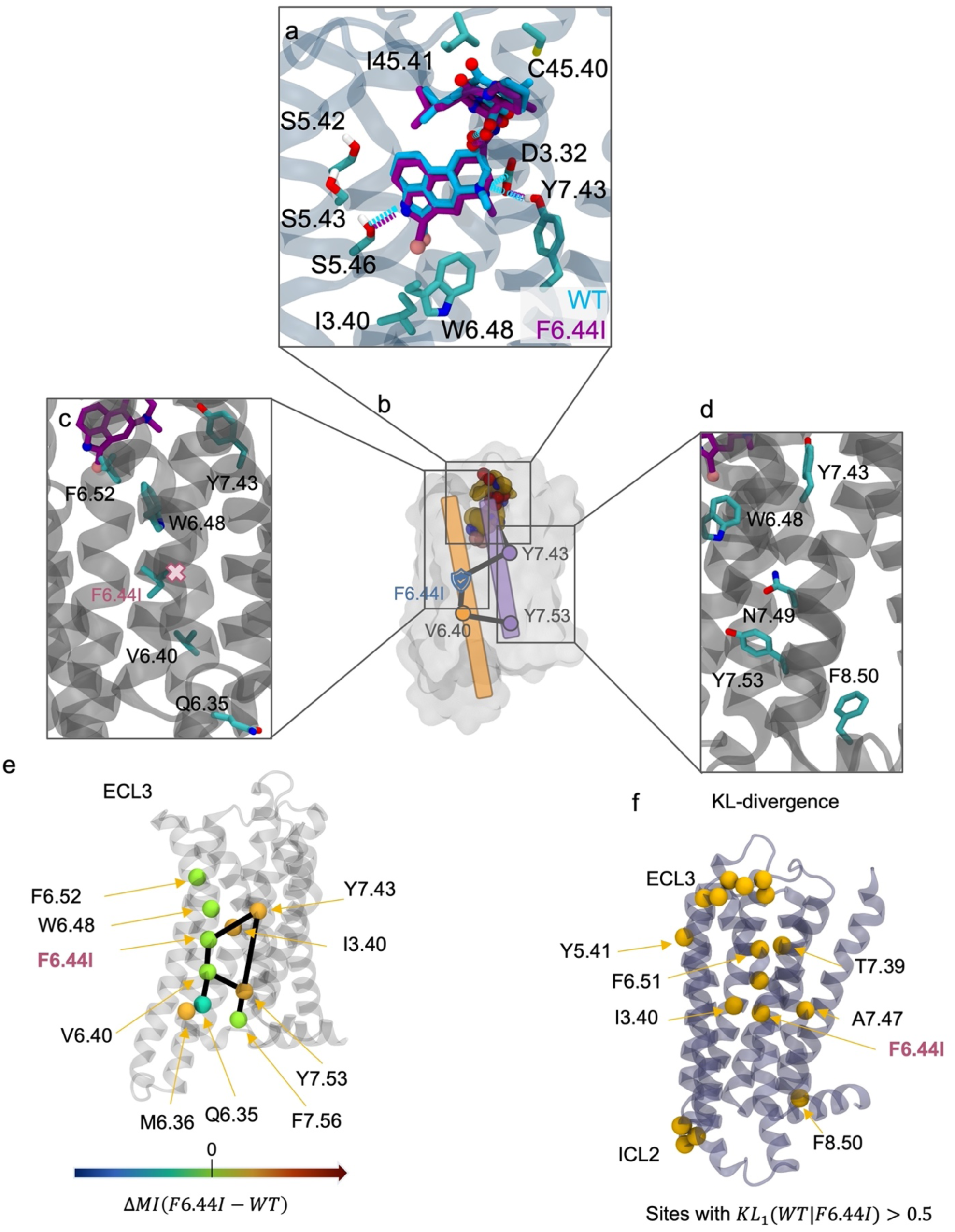
F6.44I impact on D2 allosteric response to BRC. **a.** Representative frame of highest density ligand binding pose for WT (cyan) and F6.44I (purple) extracted using PCA. **b.** Effect of the mutation as an allosteric transmitter **c.** Zoomed-in view of helix 6 showing the residues transmitting the signal through the mutation site **d.** Closeup of helix 7 showing residues involved in path propagation. **e.** Sites with large differences in mutual information (MI) mapped on the DD2R structure. MI is calculated as a sum between a given residue and all other residues. The major path connection differences from WT are represented using straight black lines and broken paths using T lines. **f.** KL-divergence visualized on the structure for residues having KL_1_ > 0.5 **Supplementary note for panel f.** Analysis of the calculated allosteric perturbations identifies the following major impacts of F6.44I on D2 allosteric responses to BRC: 1. I6.44 significantly affects the distribution of dihedrals of I3.40, more so than M6.44 2. Y7.43 establishes new allosteric connections directly with I6.44. 3. Y7.43 communicates with NPxxY and G-p binding interface through both TM6 (I6.44 and V6.40 to Q6.35 and M6.36) and TM7 4. There is comparable communication in TM5 between the mutant and WT 5. There is significant allosteric coupling to 7.53 from both TM6 and TM7 which is the hub distributing signal to the G-protein binding residues: 6.36 and 7.56. **Conclusion**: F6.44I allows allosteric signalling through the mutation position and thus restores function with BRC when compared to F6.44M.

**Supplementary Figure 21.**
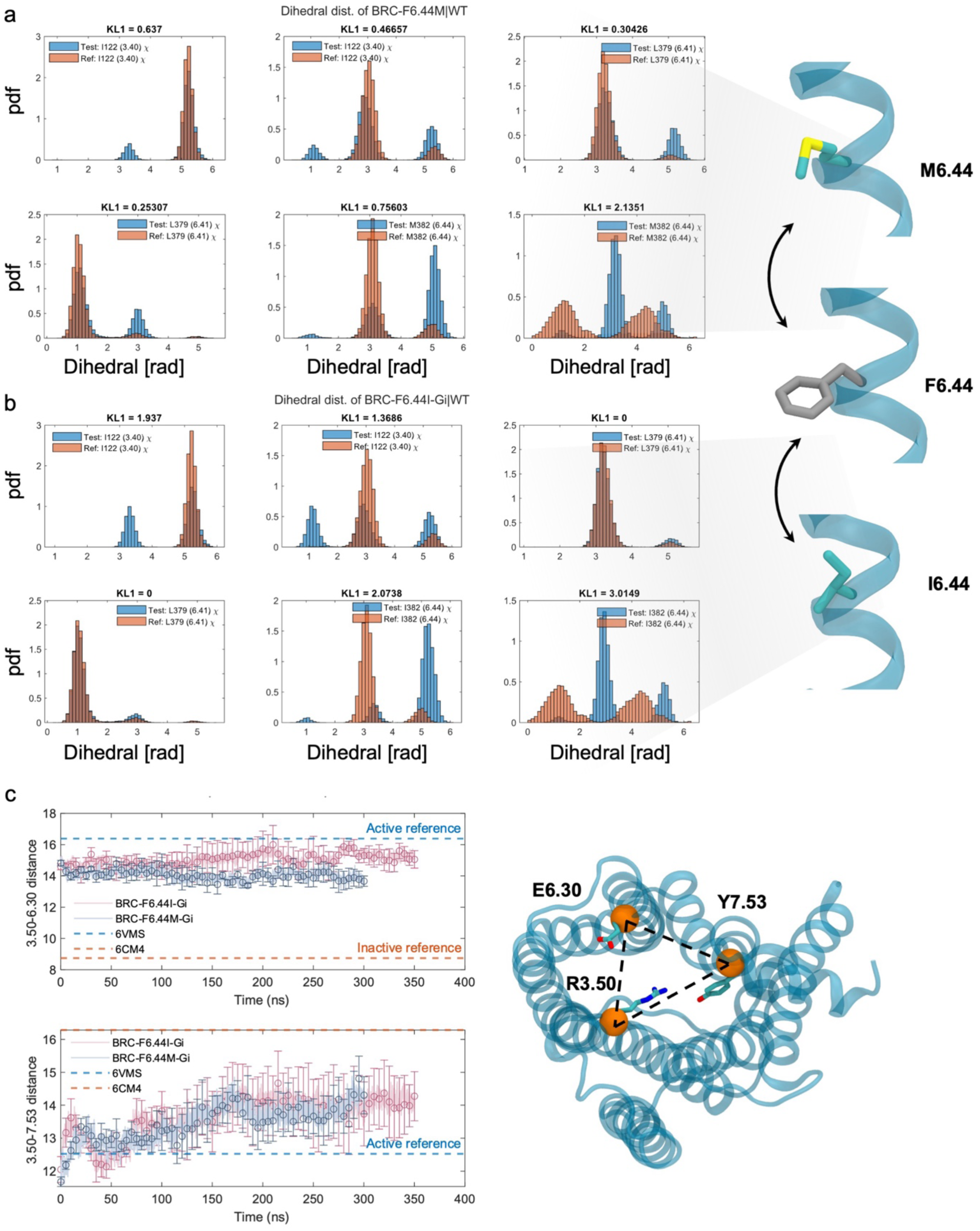
Dihedral distributions and TM distances in D2-F6.44X variants: a. and b. dihedral distributions for sites I3.40, L6.41, and X6.44 compared to WT for variants M6.44 (panel a) and I6.44 (panel b). c. TM distance distributions between TM3 (R3.50) and TM6 (E6.30) (top) and between TM3 (R3.50) and TM7 (Y7.53) (bottom) for I6.44 and M6.44 simulations. The plot shows mean values and standard deviations over 5 replicas. These distances are used as indicators of degree of activation of the receptor.

**Supplementary Figure 22:**
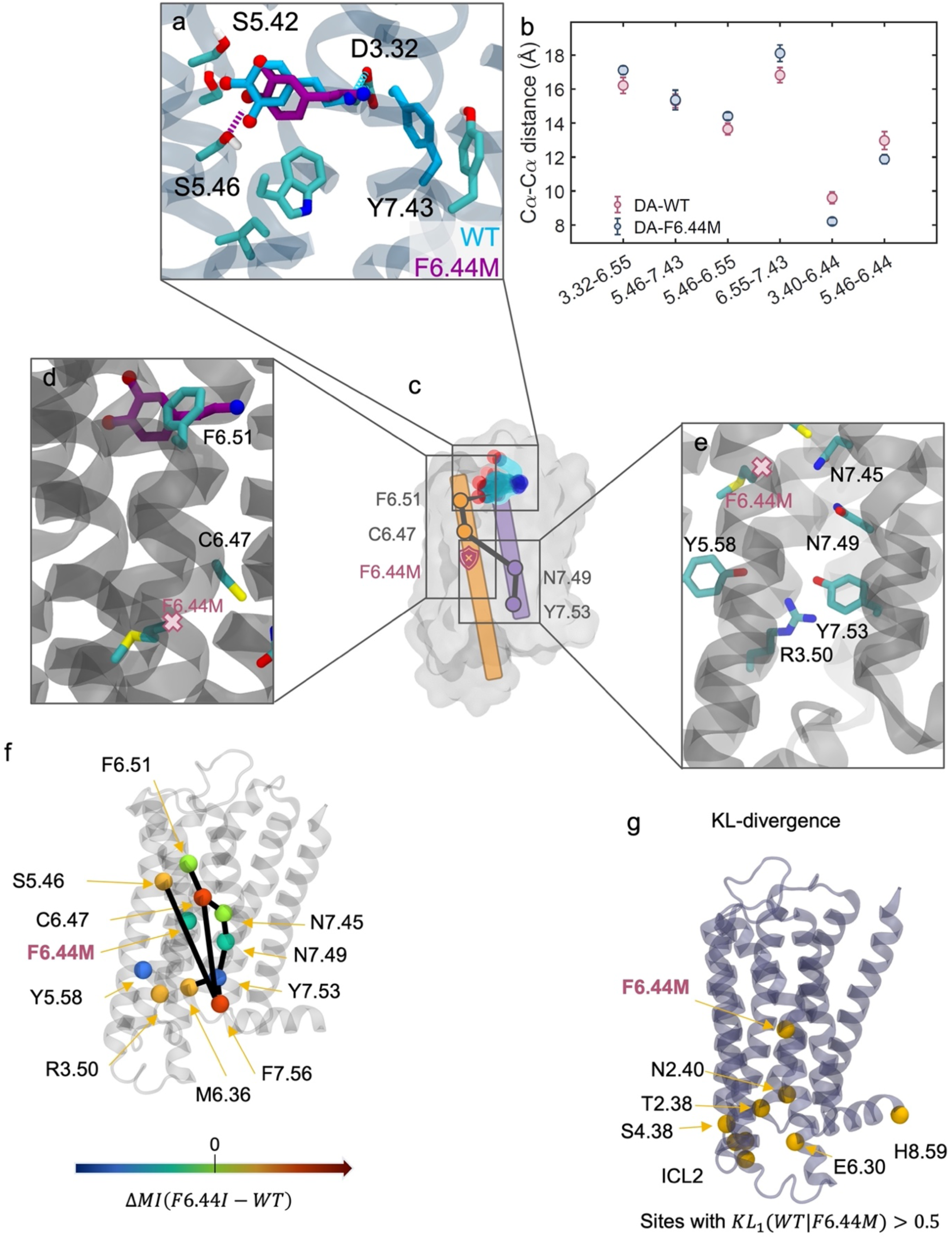
F6.44M impact on D2 allosteric response to DA. **a.** Representative frame of highest density ligand binding pose for WT (cyan) and F6.44M (purple) extracted using PCA. **b.** Distances for ligand binding residues and PIF motif layer for DA WT and F6.44M. Means and SEM are shown. Simulation frames were taken from the most populated DA binding cluster as identified by PCA (**methods**) **c.** The mutation acts as an allosteric block and triggers a conformational repositioning of DA in the binding site (**a**) that is associated with a major path rewiring along TM6 and TM7 reaching the G-protein binding surface. **d.** Zoomed in view on the mutation site and helix 6 showing the residues transmitting the signal leading to the allosteric rewiring. **e.** Zoomed in view on the mutation site and helix 7 showing the continuation of the path on conserved residues in helix 7 **f.** Sites with large differences in mutual information (MI) mapped on the DD2R structure. MI is calculated as a sum between a given residue and all other residues. The major path connection differences from WT are represented using straight black lines and broken paths using T lines. **g.** KL-divergence visualized on the structure for residues having KL_1_ > 𝟎. 𝟓. **Supplementary note for panel f.** Analysis of the calculated MI perturbations defines the following major impacts of F6.44M on D2 allosteric dynamic responses to DA: 1. F6.44M triggers is a slight loss of MI connections on TM6 around 6.44 2. However, C6.47 becomes substantially more dynamically coupled to ligand binding residues and TM7. 3. F6.44M induces significant conformational changes in the ligand binding site, shifting DA’s major binding pose. 4. DA displays stronger allosteric contacts with S5.46, F6.51, and Y7.43, enhancing communication to C6.47 and subsequently to F7.56. 5. C6.47 connects with TM7 through coupling to N7.45, propagating signals down to the NPxxY motif and the G-protein binding interface. **Conclusion**: F6.44M acts as an allosteric block with DA, but DA adapts its binding pose and circumvent the block by communication crossing from TM6 to TM7.

**Supplementary Figure 23:**
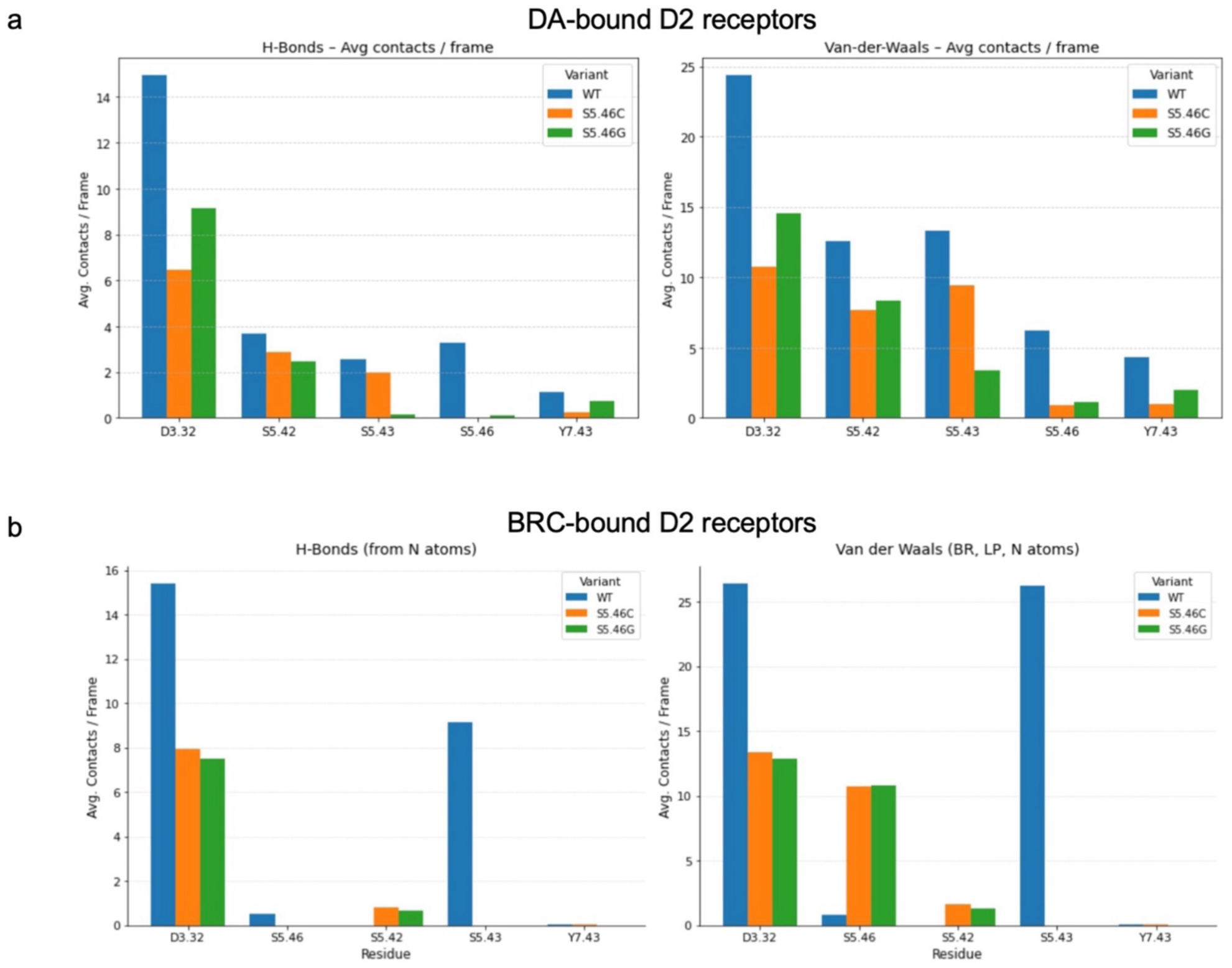
Designed mutations at position 5.46 impact ligand binding differently. Left. Average hydrogen-bond mediated contacts per frame along MD simulations of Dopamine-bound (a) and Bromocriptine-bound (b) D2 variants. Right. Average Van-der-Waals contacts per frame along MD simulations of Dopamine-bound (a) and Bromocriptine-bound (b) D2 variants. Contacts are reported for the WT (blue), S5.46C (orange) and S5.46G (green) receptor complexes.

**Supplementary Figure 24:**
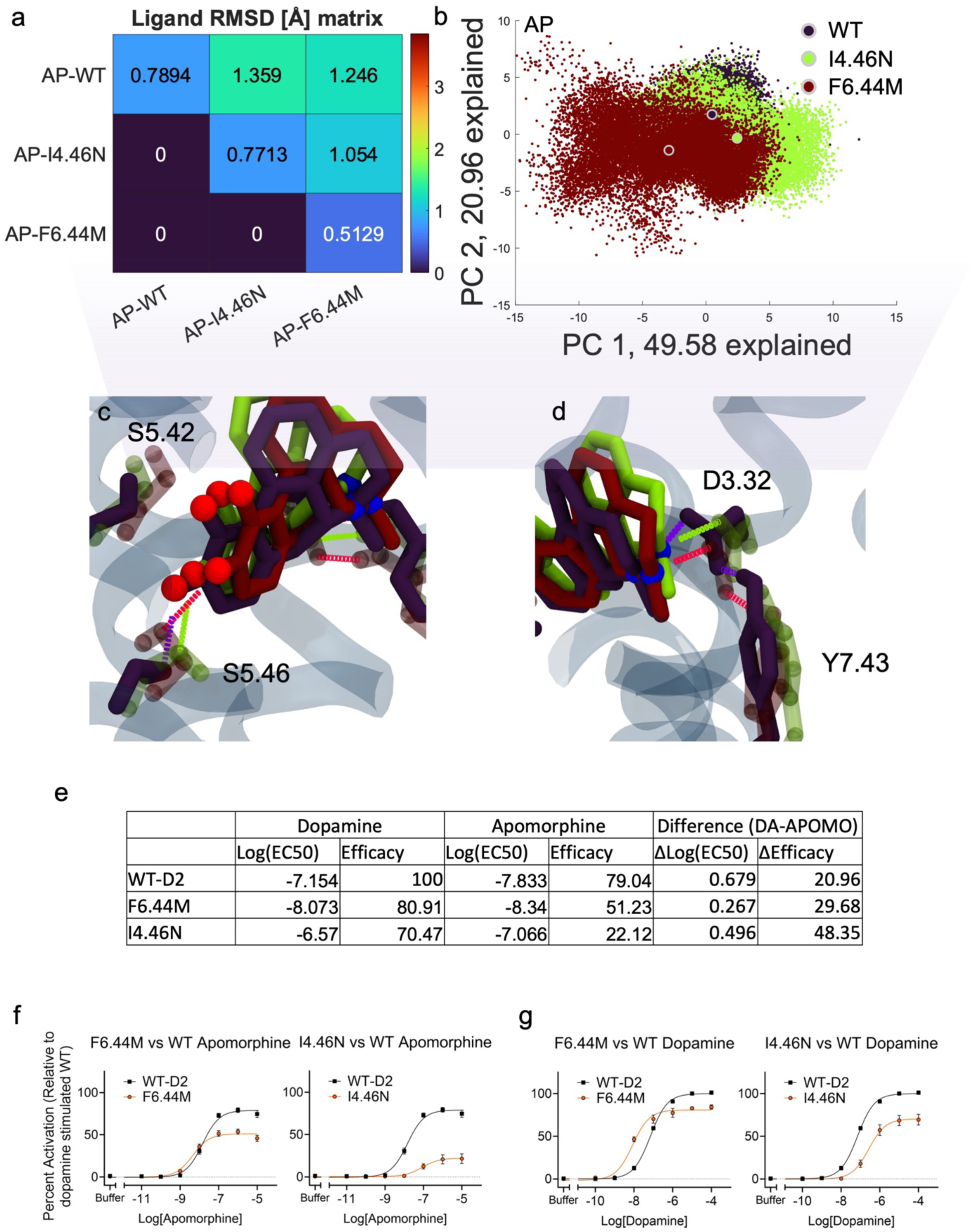
Impact of I4.46N and F6.44M on D2 allosteric response to apomorphine. Principal component analysis of the ligand binding poses of apomorphine during molecular dynamics simulations for the following variants: WT, F6.44M, and I4.46N. **a.** ligand RMSD matrix for the studied variants. RMSD is calculated by first finding highest density points for every system in PC space via histogramming (using 51 bins in every dimension). The 10 frames that are closest to the highest density PC point are then used to calculate RMSD. The mean is shown in the matrix. **b.** PC scatter for the 2 first principal components of ligand heavy atoms. The labelled dots represent the centroid of the PC spread for a given variant. The percentage of variance of the data explained by every PC is in the axis labels. **c.** and **d.** Ligand pose of representative frames of the highest density bin in PC space for WT (navy), I4.46N (lime), and F6.44M (crimson) with closeup of hydrogen bond interactions between AP and ligand binding residues **e.** Table showing ligand efficacies and potencies for DA and AP for WT and mutants **f.** and **g.** Dose response curves for apomorphine (**f.**) and dopamine (**g.**) for I4.46N (right) and F6.44M (left).

**Supplementary Figure 25.**
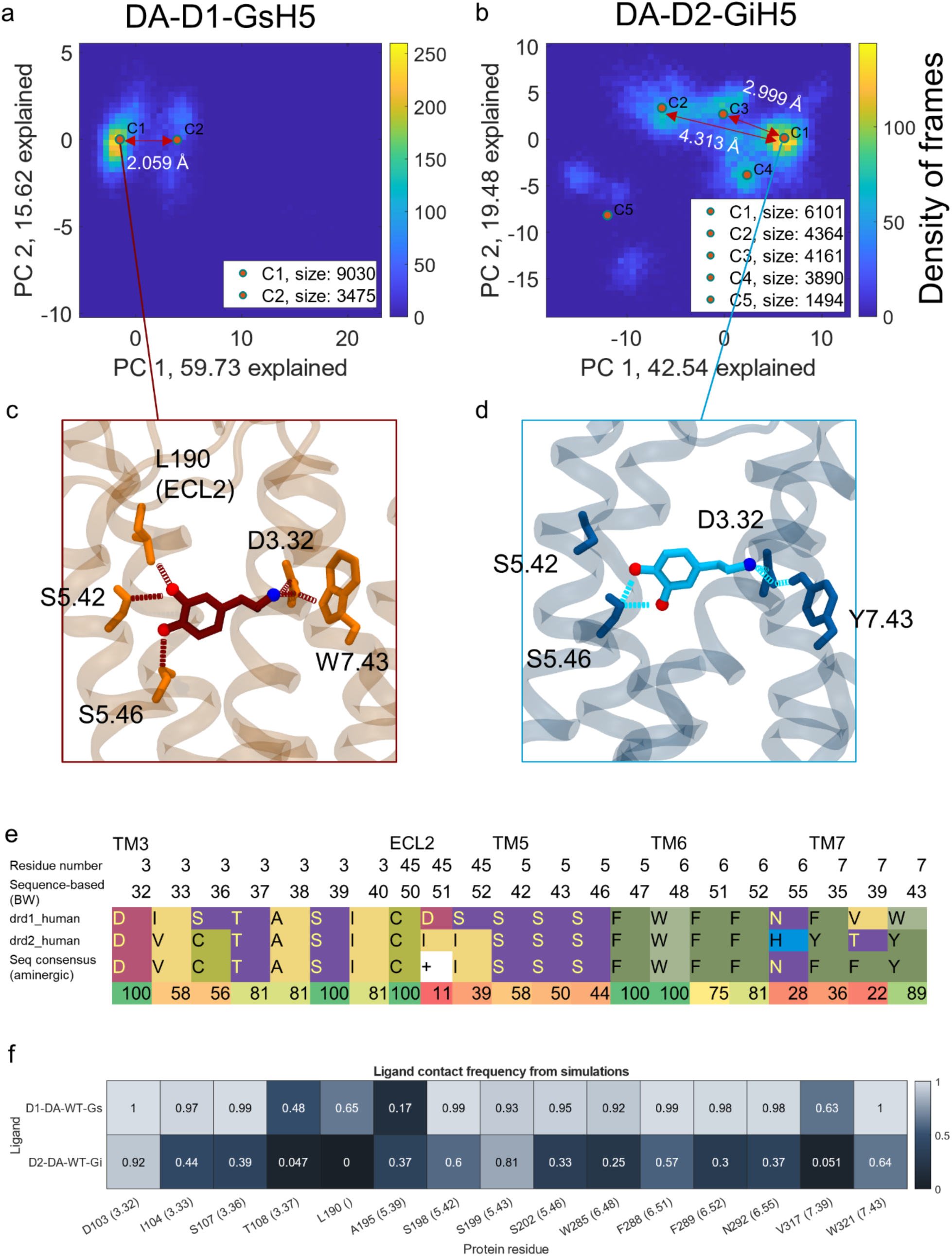
Ligand binding pose analysis for DA-bound WT D1 and D2: a, **b.** PCA was performed on ligand heavy atoms extracted from DA-D1-GsH5 (**a**) and DA-D2-GiH5 simulations (**b**). Data was clustered with 2 PCs using a K-means algorithm. **c, d.** Ligand pose of representative frames of the highest density bin in PC space for D1 (**c**) and D2 (**d**). **e.** Sequence conservation for the ligand binding residues for D1 and D2. Sequence consensus among aminergic GPCRs is shown. **f.** Ligand contacts from MD simulations for dopamine. Contacts are represented as fraction of total simulation time.

**Supplementary Figure 26:**
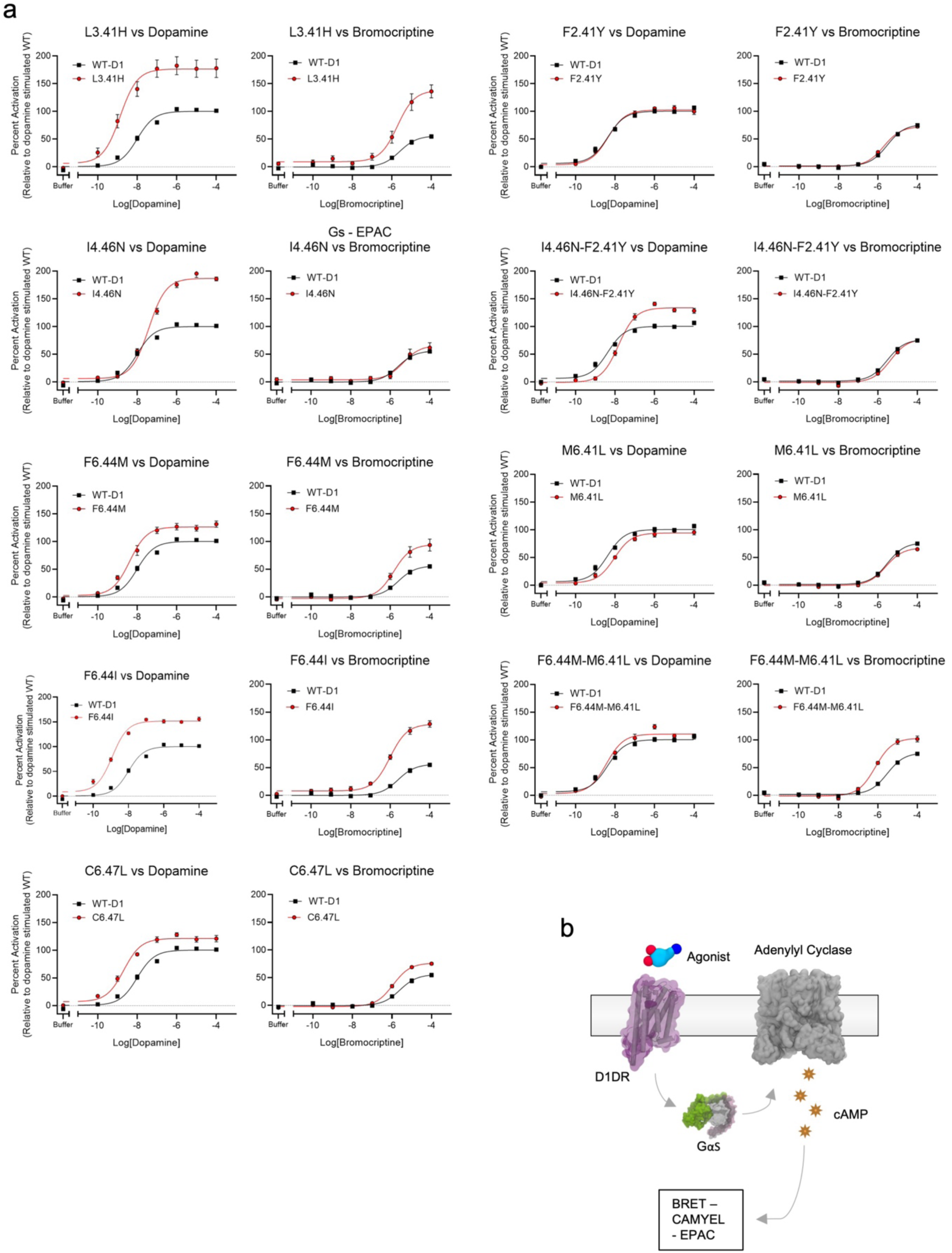
a. Experimental characterization of D1 variants. **a**. Dose Responses of D1 variants using the CAMYEL-BRET sensor system with dopamine and bromocriptine, (n=3 independent experiments made in triplicates, SEM). b. Schematic of the dopamine D1 receptor experimental system used in the dose responses in a.

**Supplementary figure 27:**
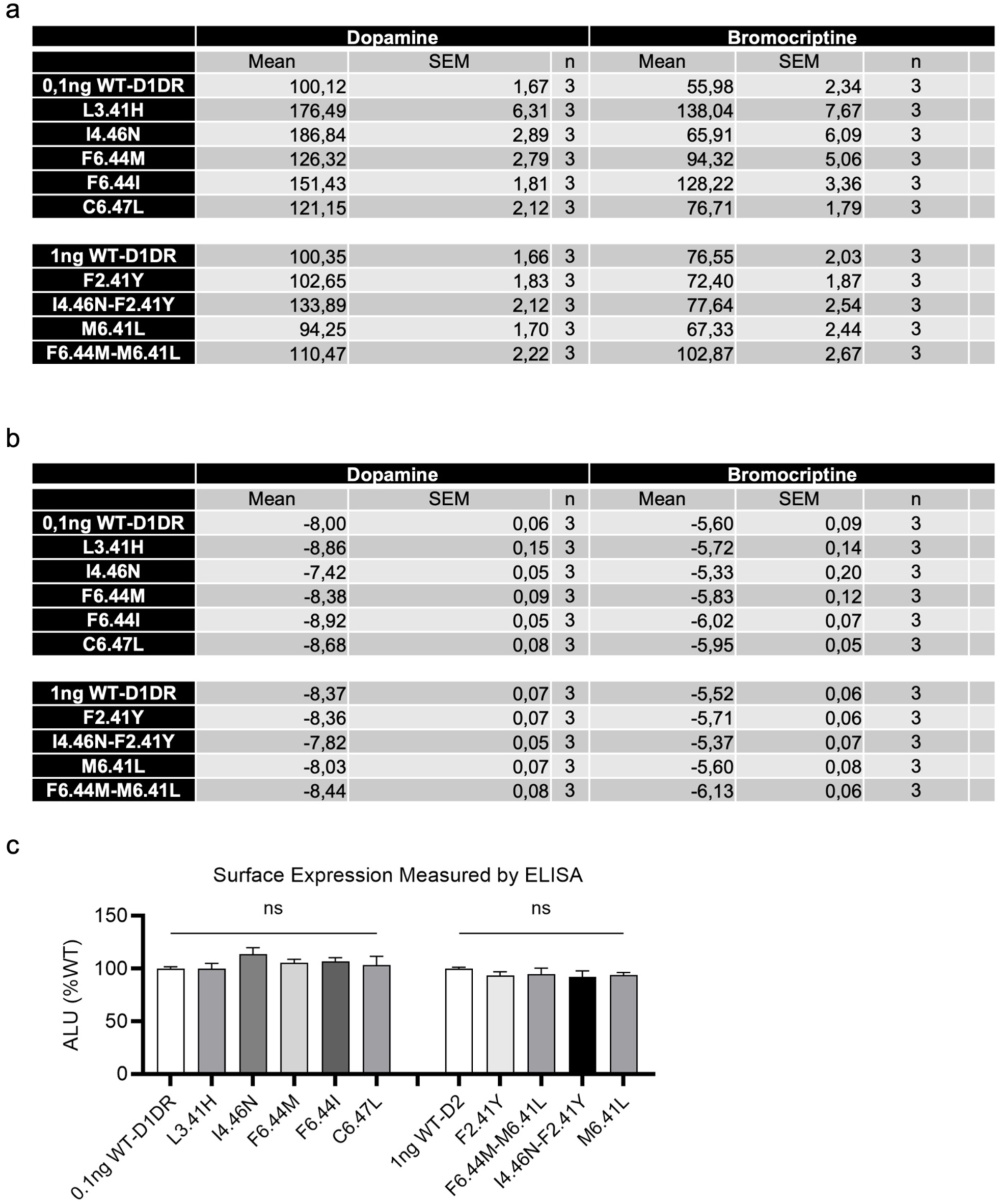
Ligand efficacy and potency for D1 variants. **a.** Mean log(EC_50_) and **b.** mean efficacies of selected mutants of dopamine D1 receptor. Standard error of the mean shown. n is the number of independent experiments, all made in triplicates. F2.41Y, I4.46N- F2.41Y, M6.41L and F6.44M-M6.41L were done with 1ng/well WT-D1DR as reference, all other with 0,1ng/well WT-D1DR. **c.** Surface expression of selected mutants of dopamine D1 receptor compared to WT-D1. Standard error of the mean shown. N=3 independent experiments, each made in triplicates

**Supplementary Figure 28:**
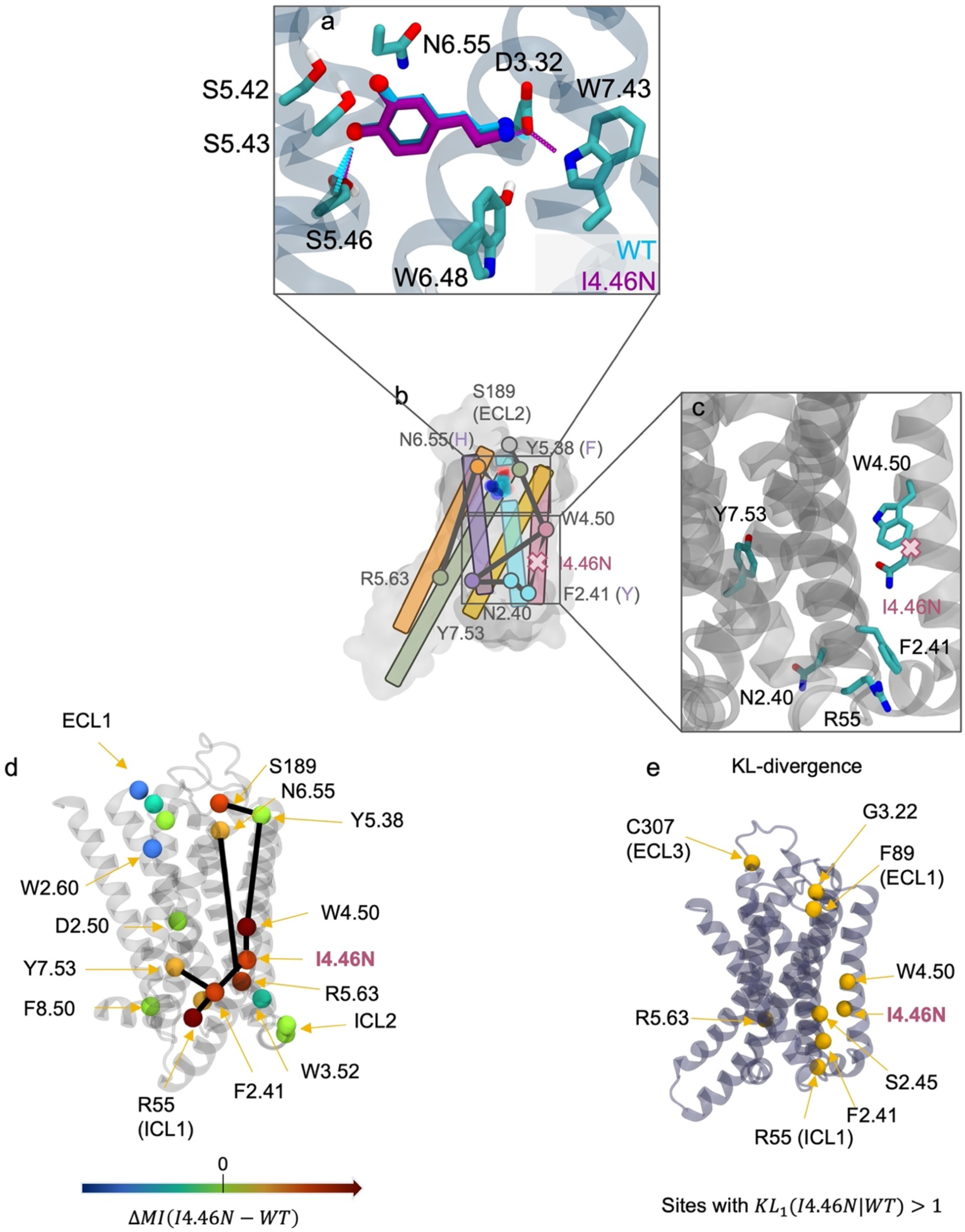
I4.46N impact on D1 allosteric response to DA. **a.** Representative frame of highest density ligand binding pose for WT (cyan) and I4.46N (purple) extracted using PCA. **b.** The mutation acts as an allosteric enhancer. **c.** Zoomed in view on the mutation site and helix 7 showing the local interactions of the mutated site **d.** Sites with large differences in mutual information (MI) mapped on the DD1R structure. MI is calculated as a sum between a given residue and all other residues. The major path connection differences from WT are represented using straight black lines and broken paths using T lines. **e.** KL-divergence visualized on the structure for residues having KL_1_ > 𝟏 **Supplementary note for panel e.** Analysis of the calculated MI perturbations and KL-divergences defines the following major impacts of I4.46N on D1 allosteric dynamic responses to DA: 1. I4.46N impacts the conformation of neighboring residues (F2.41, S2.45, and W4.50) through multiple polar and hydrophobic contacts. 2. Significant increase in MI is observed for several residues connecting the extracellular to the intracellular surfaces (S189, N6.55, W4.50, N4.46, N2.40, F2.41, R55, R5.63 and Y7.53) compared to WT. These strengthen allosteric pathways along TM2 and TM4 reaching the G-protein binding site. 3. DA displays stronger allosteric contacts with F6.51 and N6.55 and is more dynamically coupled to the designed receptor than WT. **Conclusion**: I4.46N acts as an allosteric enhancer with DA, with strong communication from the ligand to the G-protein binding surface passing through TM2 and TM4 and the mutated site.

**Supplementary Figure 29:**
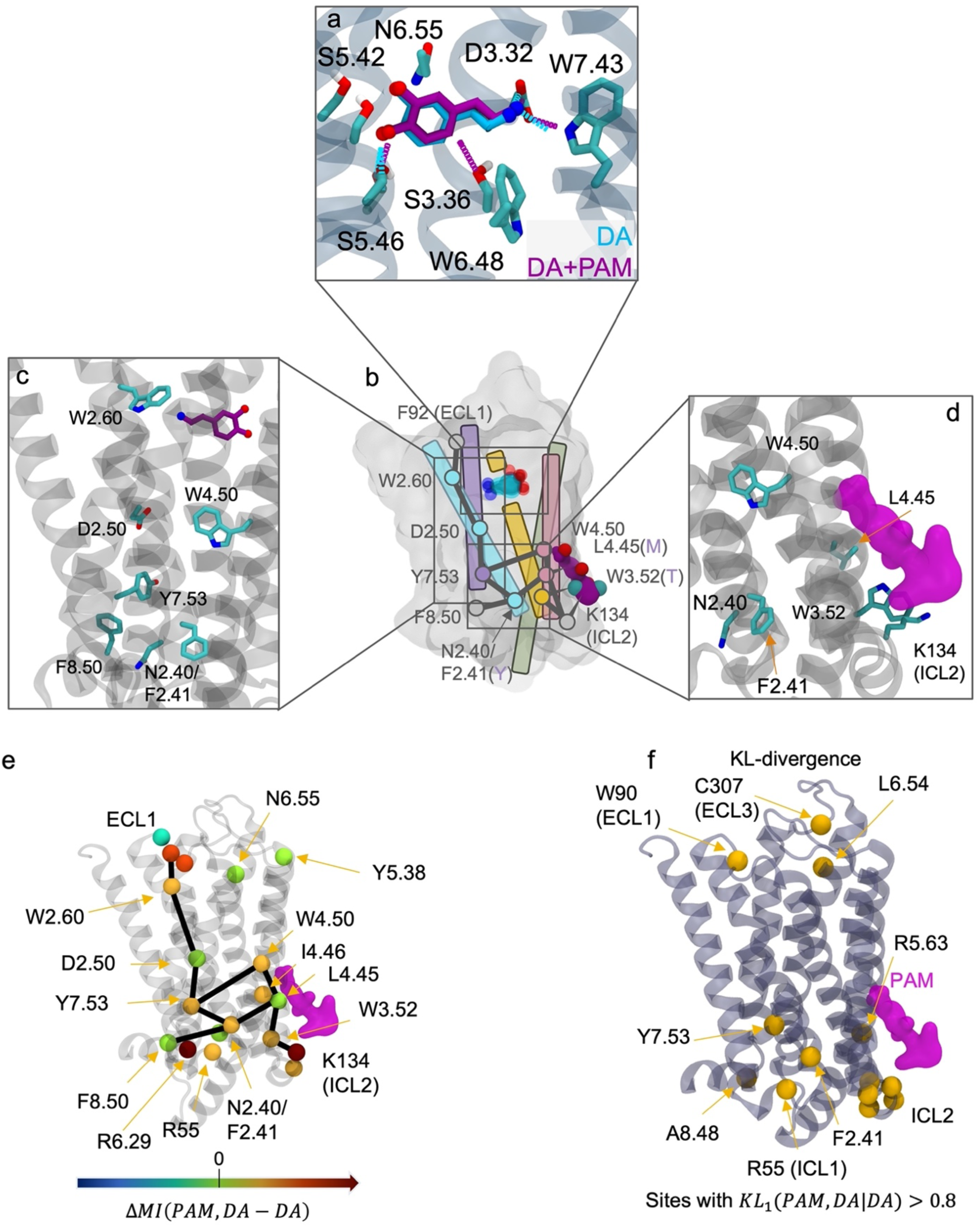
Positive allosteric modulator (PAM) binding impact on D1 allosteric response to DA. **a.** Representative frame of highest density ligand binding pose for DA only (cyan) and DA+PAM (purple) extracted using PCA. **b.** Effect of PAM as an allosteric enhancer. **c.** Zoomed in view on helix 7 showing the residues transmitting the signal leading to the allosteric enhancement **d.** Zoomed in view on the PAM binding site showing local interactions **e.** Sites with large differences in mutual information (MI) mapped on the DD1R structure. MI is calculated as a sum between a given residue and all other residues. The major path connection differences from WT are represented using straight black lines and broken paths using T lines. **f.** KL-divergence visualized on the structure for residues having KL_#_ > 𝟎, 𝟖. **Supplementary note for panel f.** Analysis of the calculated MI perturbations and KL-divergences defines the following major impacts of PAM-binding on D1 allosteric dynamic responses to DA: 1. The PAM binding triggers numerous long-range allosteric structural perturbations in the extracellular and intracellular binding regions in addition to side-chain conformational changes in the direct vicinity of the PAM on TM4 and ICL2 (**g**). 2. We also observed a significant increase in total MI across the entire receptor (**f**) but especially at residues K134 (ICL2), I4.46, W4.50, R6.29, and Y7.53. 3. These shifts in dynamic coupling rewire the allosteric pathways, connecting ECL1 and PAM binding site to NPxxY motif and the G-protein binding interface. **Conclusion**: PAM binding acts as an allosteric enhancer for DA, increasing communication from DA through the PAM binding site to Y7.53 and multiple residues on the G-protein binding surface.

**Supplementary Figure 30:**
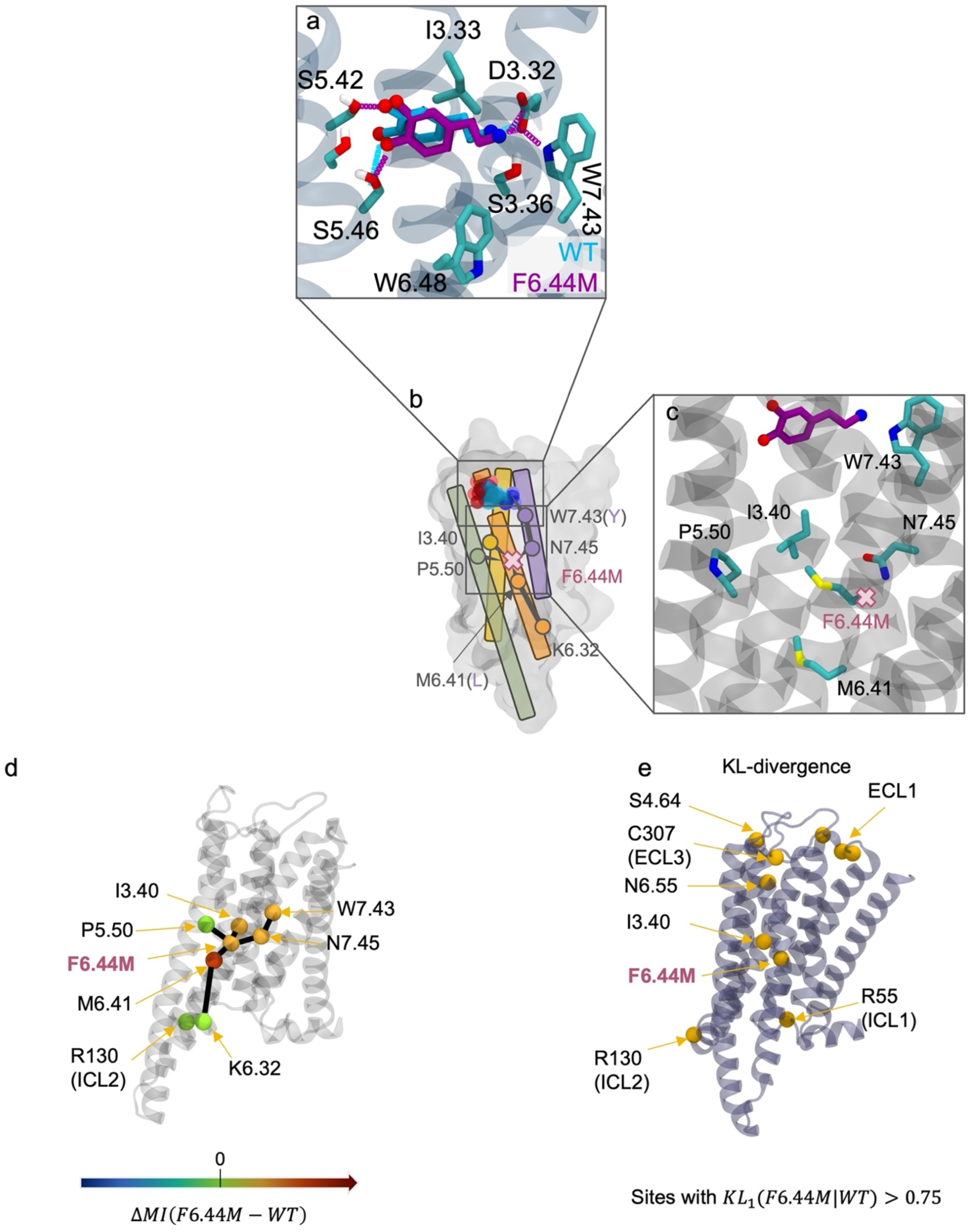
F6.44M impact on D1 allosteric response to DA. **a.** Representative frame of highest density ligand binding pose for WT (cyan) and F6.44M (purple) extracted using PCA. **b.** Effect of mutation as an Allosteric transmitter **c.** Zoomed-in view on helices 5 and 6 showing the residues transmitting the signal **d.** Sites with large differences in mutual information (MI) mapped on the D1 structure. MI is calculated as a sum between a given residue and all other residues. The major path connection differences from WT are represented using straight black lines and broken paths using T lines. **e.** KL-divergence visualized on the structure for residues having KL_1_ > 𝟎, 𝟕𝟓. **Supplementary note for panel e.** Analysis of the calculated MI perturbations and KL-divergences defines the following major impacts of F6.44M on D1 allosteric dynamic responses to DA: 1. We observe a significant increase in dynamic communication along TM6 and 7 (especially at residues M6.41, M6.44, N6.55, W7.43, and Y7.53). 2. Allosteric contacts between DA and residues on TM6 and 7 became also significantly stronger than with WT D1. 3. Allosteric communication paths are initiated from the ligand binding contacts at TM7 and TM6 reaching position 6.44. They continue along TM6 thanks to strong dynamic coupling between M6.44 and M6.41 down to the G-protein binding region. **Conclusion**: Unlike in D2, F6.44 acts as an allosteric transmitter for DA, and enables communication from the ligand binding region through the mutated site down to the G-protein binding interface.

